# Development of a Nanoparticle-Based Tendon-Targeting Drug Delivery System to Pharmacologically Modulate Tendon Healing

**DOI:** 10.1101/2023.11.29.569204

**Authors:** Emmanuela Adjei-Sowah, Indika Chandrasiri, Baixue Xiao, Yuxuan Liu, Jessica E. Ackerman, Celia Soto, Anne E. C. Nichols, Katherine Nolan, Danielle S.W. Benoit, Alayna E. Loiselle

## Abstract

Tendon regeneration following acute injury is marred by a fibrotic healing response that prevents complete functional recovery. Despite the high frequency of tendon injuries and the poor outcomes, including functional deficits and elevated risk of re-injury, there are currently no pharmacological therapies in clinical use to enhance the healing process. Several promising pharmacotherapies have been identified; however, systemic treatments lack tendon specificity, resulting in poor tendon biodistribution and perhaps explaining the largely limited beneficial effects of these treatments on the tendon healing process. To address this major unmet need, we leveraged our existing spatial transcriptomics dataset of the tendon healing process to identify an area of the healing tendon that is enriched for expression of *Acp5. Acp5* encodes tartrate-resistant acid phosphatase (TRAP), and we demonstrate robust TRAP activity in the healing tendon. This unexpected finding allowed us to refine and apply our existing TRAP binding peptide (TBP) functionalized nanoparticle (NP) drug delivery system (DDS) to facilitate improved delivery of systemic treatments to the healing tendon. To demonstrate the translational potential of this drug delivery system, we delivered the S100a4 inhibitor, Niclosamide to the healing tendon. We have previously shown that genetic knockdown of S100a4 enhances tendon healing. While systemic delivery of Niclosamide did not affect the healing process, relative to controls, TBP-NP delivery of Niclosamide enhanced both functional and mechanical outcome measures. Collectively, these data identify a novel tendon-targeting drug delivery system and demonstrate the translational potential of this approach to enhance the tendon healing process.

## Introduction

Tendons are dense, connective tissues that transmit forces from muscle to bone. After an injury, tendons heal in a fibrotic manner, characterized by the deposition of excessive and disorganized extracellular matrix ^1, 2^. The current gold standard for treating acute tendon injuries is surgical intervention, however, poor outcomes, including re-rupture, and peritendinous fibrosis are common ^3^. Given that there are over 300,000 surgical tendon repairs in the United States annually ^1, 2, 4^, unsatisfactory outcomes following tendon injuries represent a major clinical burden, and there is a clear need for biological strategies to enhance the healing process. Furthermore, tendons do not typically regain native mechanical and functional properties following injury ^1, 2, 4^. However, very few pharmacological approaches to enhance tendon healing have been identified, and systemic treatments are plagued by poor tendon-targeting. As such, newly developed treatment modalities must address these two unmet needs.

S100a4 is a calcium-binding protein implicated in fibrosis in many tissues, including the liver ^5^, heart ^6^, lung ^7^, oral submucosa ^8^, and, recently, the tendon ^9–11^. Previous studies from our lab utilizing a genetic mouse model of S100a4 haploinsufficiency have demonstrated the benefits of S100a4 knockdown during scar-mediated tendon healing, highlighting S100a4 as a potent, anti-fibrotic therapeutic target to improve tendon healing ^11^. While genetic mouse models are an important tool for candidate identification, direct clinical translation of therapeutic interventions based on these studies is very limited. However, FDA-approved small molecule S100a4 inhibitors may represent greater translational potential to improve tendon healing.

Local administration of therapeutics to the tendon is the most common approach for tendon-specific drug delivery, as systemic delivery is limited by poor tendon vascularization ^12^. Nevertheless, local delivery has several limitations, including short tissue half-life due to diffusion, and tissue damage due to injection and/or locally high initial drug concentrations^12, 13^. While local drug delivery systems can overcome some of these limitations, they must maintain controlled release capabilities in the dynamic biomechanical environment ^12–15^. Based on the limitations of local administration, systemic delivery is appealing. However, small molecules have low bioavailability, with less than 1% of systemically administered drugs reaching the tendon ^14, 15^. Poor tendon specificity requires concomitantly high doses of drug to reach therapeutic efficacy, resulting in off-target effects ^12^. Drug delivery systems (DDS), including nanoparticles, can be used to increase drug solubility and bioavailability. However, nanoparticles have very low site-specific targeting, and must, therefore, be functionalized with targeting ligands ^16, 17^. Unfortunately, no strategies to target the tendon have been identified.

Hence, our overarching goal was to identify tendon targeting moieties to enable efficient, tendon-specific drug delivery. In this study, a poly(styrene-alt-maleic-anhydride) (PSMA-*b*-PS) NP delivery system, which offers extensive chemical versatility for functionalization with various ligands for tissue-specific targeting was used. To enhance tendon-specific targeting of these NPs, we leveraged our prior spatial transcriptomic dataset to identify novel molecular markers for targeting^18^. We discovered that the tendon healing site was enriched in expression of *Acp5*, the gene encoding Tartrate Resistant Acid Phosphatase (TRAP). While TRAP is typically associated with bone-remodeling osteoclasts, high TRAP activity within the healing tendon allows us to leverage a peptide with subnanomolar affinity for TRAP^19^. Therefore, we hypothesize that conjugation of the TRAP binding peptide to the PSMA-*b*-PS NPs facilitates high-efficiency targeting of the healing tendon and, in combination with delivery of an S100a4 inhibitor, develops a novel tendon drug delivery system with promising translational potential.

## Results

### TRAP is expressed in the injured tendon

The fundamental cellular and molecular mechanisms that drive scar-mediated tendon healing are poorly defined. Therefore, we utilized the 10X Spatial Transcriptomics platform to comprehensively define the predominant spatial-molecular programs during tendon healing ^18^. Integrated analysis of data from uninjured tendons, and D14, D21, and D28 post-repair, identified five distinct spatiomolecular clusters in the tendon (**Fig. 1A**)^18^, including spatially overlapping clusters at the healing site that were annotated as ‘inflammatory/immune’ (cluster 4) and ‘reactive tissue’ (cluster 2) (**Fig. 1A**). The immune cluster was enriched for macrophage markers including *Mmp9* and *Mmp13*, however the most differentially expressed gene in this cluster was *Acp5*, the gene encoding for TRAP (**Fig. 1B**). Spatial mapping of *Acp5* expression at D14 demonstrated robust *Acp5* expression in both the tendon stubs and bridging scar tissue (**Fig. 1C**). While TRAP is typically associated with bone-remodeling osteoclasts, other cells in the hematopoietic lineage also express TRAP ^20^. Based on the surprising finding of high *Acp5* expression in the healing tendon, we performed staining for TRAP activity during tendon healing. No TRAP activity was observed in the uninjured tendon (**Fig. S1A**), and minimal TRAP activity was observed before D7 post-surgery (**Fig. S1B**). By D7, TRAP+ cells were observed in the bridging granulation tissue between the tendon stubs (**Fig. 1D**). At D14 a substantial increase in TRAP staining was observed, with TRAP activity observed diffusely throughout the native tendon stubs (**Fig. 1D**), and in the bridging tissue between the tendon stubs (**Fig. 1D**). Collectively, these data demonstrate robust TRAP activity in the healing tendon, supporting the potential adoption of our previously reported TRAP Binding Peptide nanoparticle (TBP-NP) system^19^ for high efficiency drug delivery to the healing tendon.

**Figure 1.**
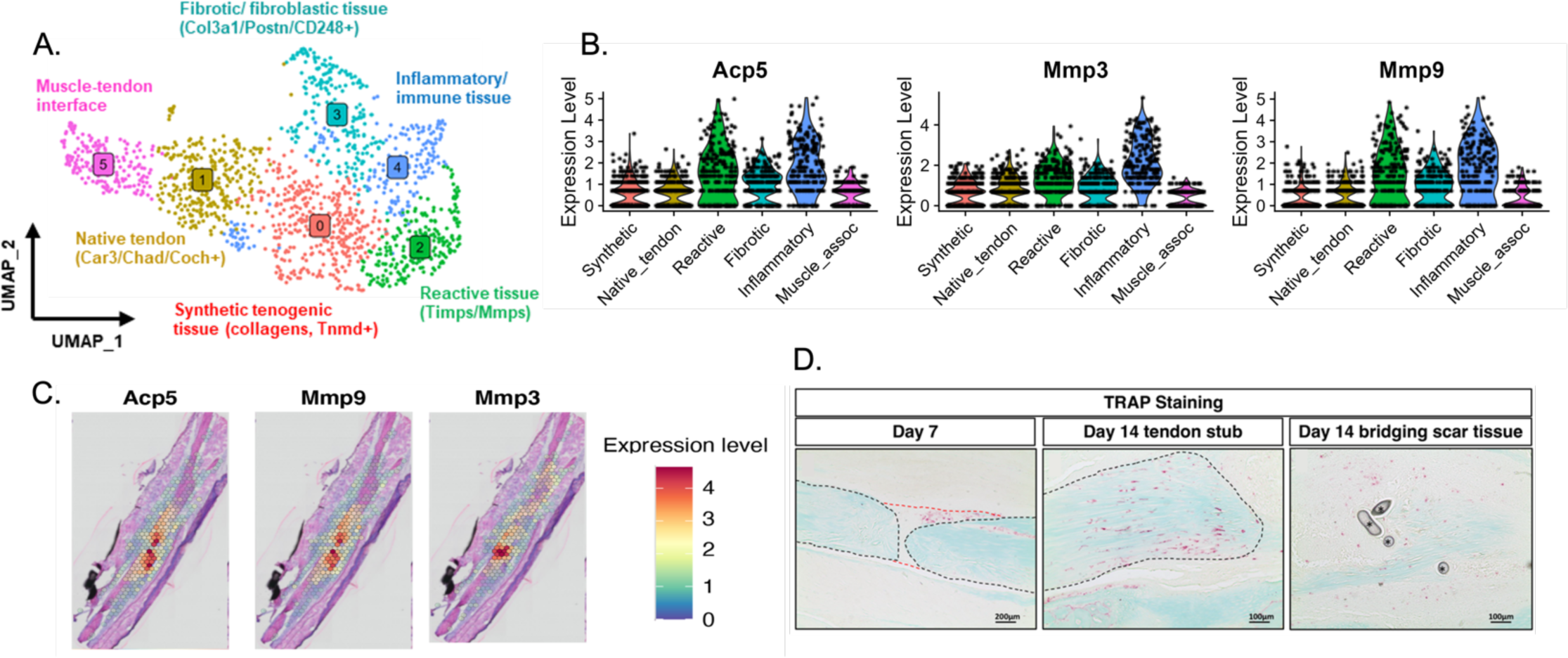
Spatial transcriptomic analysis of tendon healing identifies an inflammatory/macrophage cluster at the tendon repair site. (A) UMAP analysis of unsupervised clustering of spatial transcriptomics data from uninjured tendons and tendons at D14 and 28 days post-repair, identifies 5 distinct molecular clusters ^21^. (B) Cluster 4 which is defined as an inflammatory cluster is defined in part by high expression of Acp5. (C) Mapping of *Acp5* demonstrates high expression and specific localization in the tendon stubs and bridging tissue. (D) High levels of TRAP activity (red) are observed in the healing tendon. At D7, a cluster of TRAP cells is observed in the bridging tissue, with an additional TRAP+ population in tendon stub (outlined in black). By D14 the TRAP+ population has expanded with diffuse localization throughout both the tendon stub and bridging scar tissue.

### Peptide functionalized nanoparticles are formed via RAFT polymerization and Anhydride Ring Opening reaction

While our previous use of the TBP-NP system facilitated high-efficiency targeting of fractured bone, we sought to gain greater control and reproducibility of the TBP conjugation step due to the presence of multiple amine and carboxylates. Hence, to minimize the occurrence of undesired side reactions, we moved from carbodiimide chemistry to a maleic anhydride ring opening reaction^22^. To synthesize TBP-NPs, we first used RAFT polymerization to synthesize amphiphilic diblock copolymers composed Maleic anhydride (MA) and Styrene (STY). RAFT is a favorable approach to synthesize diblock polymers because it results in polymers with low polydispersity ^23^. We synthesized amphiphilic diblock polymers by adding styrene in excess of MA in the presence of the RAFT chain transfer agent, DCT (**Fig 2A**). Thus, after preferential reactivity results in alternating copolymer blocks of styrene and maleic anhydride, a homopolymer styrene second block will result in a 1-step reaction ^24^. To characterize our polymers, GPC and H^1^NMR were used. Diblock molecular weights were of 41 ± 2.3 kDa [1^st^ block = 21 kDa, 2^nd^ block = 20 kDa], with polydispersity of 1.1 ± 0.03. Diblock copolymers were then functionalized with either a TRAP Binding Peptide (TBP) or a Scrambled Control Peptide (SCP) (**Fig 2B**). TBP is a peptide identified through phage display with sub-nanomolar affinity for TRAP ^19, 25^. SCP is a control peptide that contains the same amino acids as TBP, but is randomized. TBP and SCP conjugation to PSMA-*b*-PS was achieved through maleic anhydride ring opening reactions ^22^. Since our peptides were synthesized via solid-phase peptide synthesis on a FMOC-Gly-Wang resin, the FMOC could be cleaved without affecting the acid-labile Alloc protecting group. Alloc was deprotected using Tetrakis (Pd(PPh_3_)_4_) and Phenylsilane **(Fig. 2C)**, which acts as a scavenger through hydride donor activity ^26^. After deprotection, the peptide-polymer conjugates were purified via dialysis and characterized via NMR, which confirmed removal of allylic peaks (**Fig. S2**). Self-assembled nanoparticles (NPs) were formed from TBP- and SCP-polymer conjugates via solvent exchange from DMF to water (**Fig 2D**). We confirmed that the conjugated product had Mn of 56 ± 2.1 kDa, with a polydispersity of 1.02 ± 0.01. Polydispersity did not change appreciably from the unfunctionalized polymer, suggesting peptides were uniformly introduced to the polymer chains (**Fig. 2E**). Furthermore, the Gaussian characteristic of the PSMA-*b*-PS molecular weight distribution remained unaltered, and there were no observable additional peaks in the GPC chromatograms (**Fig. S3).** To estimate the average amount of conjugated peptides per polymer chain, we subtracted the molecular weight of the unconjugated polymer from the conjugated polymer, based on the feed ratio used. We observed close results based on feed, with an average of 9 peptides per polymer chain **(Fig. 2E).** These data indicate that we achieved a controlled conjugation reaction of the peptides and polymers.

**Figure 2.**
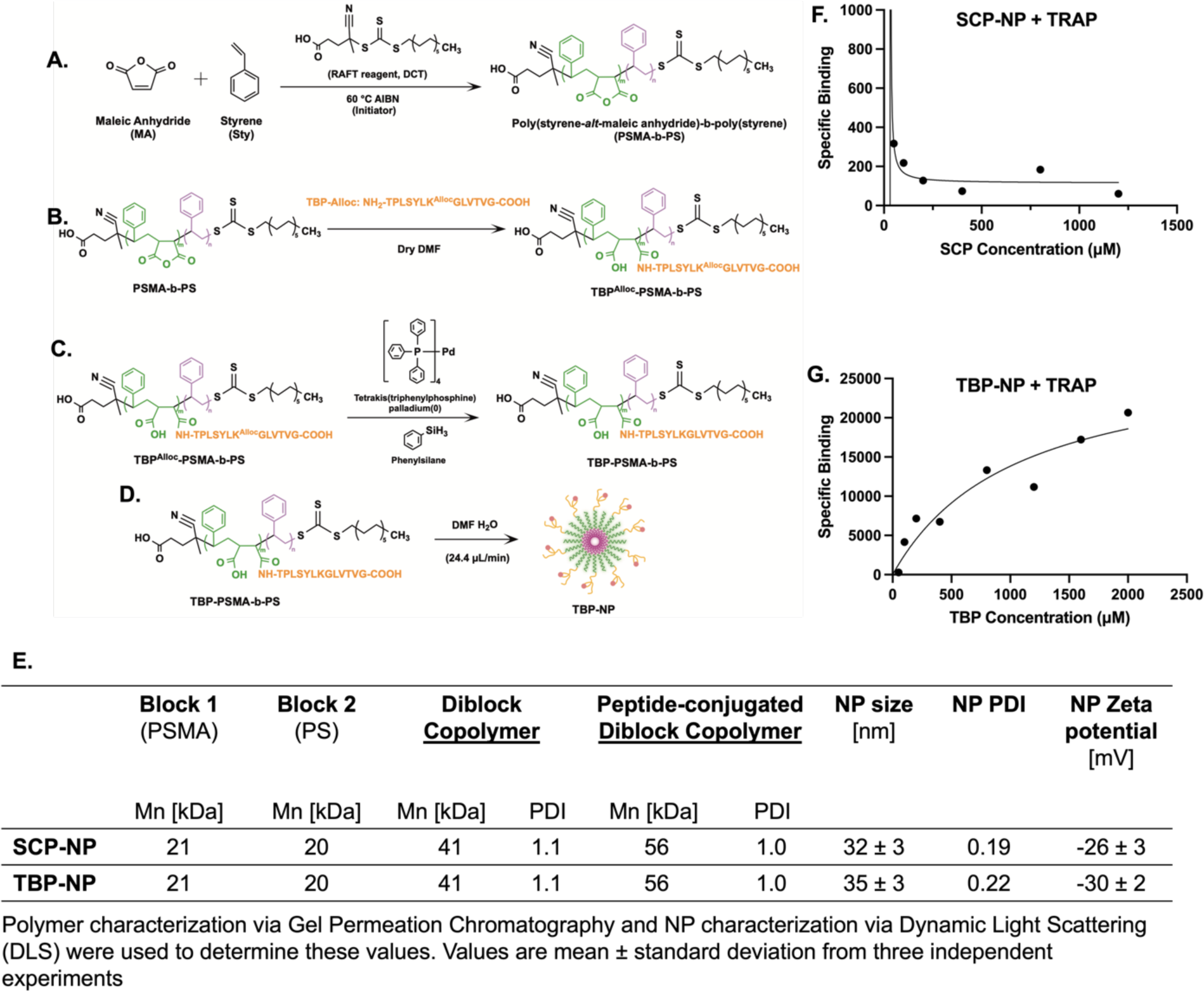
Characterization of the drug delivery system. **(A)** PSMA-b-PS polymers are formed from two monomers via RAFT. **(B)** Synthesized polymers are functionalized with a TRAP Binding Peptide (TBP) with an allylic component, **(C)** which is deprotected with Tetrakis and Phenylsilane. **(D)** Functionalized polymers are self-assembled into amphiphilic nanoparticles. **(E)** Untargeted SCP-NPs have no affinity for TRAP, whereas **(F)** TBP-NP binding behavior indicates high affinity for TRAP. **(G)** SCP-NP and TBP-NPs have desirable physicochemical properties.

TBP-NP and SCP-NP hydrodynamic diameter and surface charge were determined via DLS to be 35 ± 3 nm and 32 ± 3 nm, and −30 ± 2 mV and −26 ± 3 mV respectively (**Fig 2E).** TEM also revealed the formation of uniform NPs, consistent with NP sizes obtained from DLS (**Fig. S4).** We confirmed that TBP-NPs had a high affinity for TRAP via a TRAP binding assay. Specifically, when TBP-NPs were exposed to TRAP for 2 h, a binding affinity (K_D_) of 153 µM with a B_max_ of 258,700 was observed. SCP-NP binding curves do not have the expected behavior, therefore, are deemed to have no measurable affinity (**Fig 2F-G**). Notwithstanding, these data establish TBP-NPs as a promising drug delivery system for tendon targeting due to its attractive physicochemical properties and high affinity for TRAP.

### TBP-NPs demonstrate high tendon homing and sustained retention during tendon healing

During the acute inflammatory phase of wound healing, the Enhanced Permeability and Retention effect (EPR) allows molecules of certain sizes to accumulate in injured tissues, which can be leveraged to increase nanoparticle accumulation ^27, 28^. As such, we assessed the biodistribution of IR780-labelled TBP-NPs delivered during this period (NP administration on D3 post-surgery), relative to IR780-labelled SCP-NP treated mice and saline controls, via in vivo imaging (**Fig. 3A**). Semi-quantitative analysis of IR780 accumulation was performed by measuring the total radiant efficiency of regions of interest (right hind-paw) and normalizing to saline controls. In both TBP-NP_IR780_ and SCP-NP_IR780_ groups, accumulation of NPs peaked at ∼48-72 h post-injection (**Fig. 3B)**. We observed some accumulation of SCP-NP_IR780_ in the right hind paw, likely due to EPR-related accumulation. However, targeting via TBP showed 3-fold greater accumulation over the untargeted SCP-NP_IR780_ at D8. Furthermore, TBP-NP_IR780_ were retained in the tendon for ∼14 days, compared to SCP-NP_IR780_ which were cleared within 7 days.

**Figure 3.**
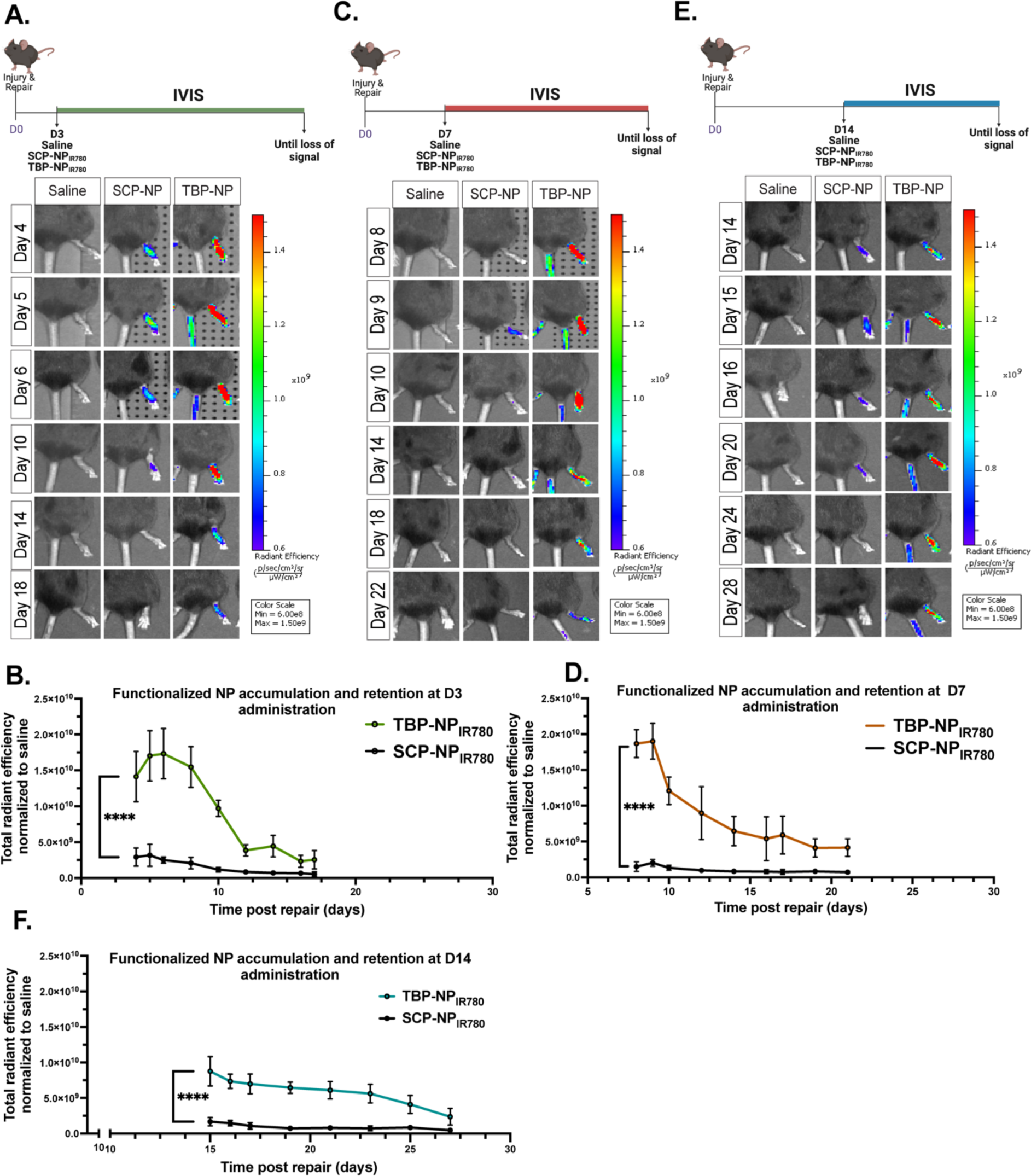
Use of a TRAP binding peptide-functionalized nanoparticle delivery system to target injured tendon. Schematic representation of treatment timeline for targeting studies. Representative live animal imaging and graphs shows biodistribution of NPs after **(A-B)** day 3 treatment **(C-D)** day 7 treatment, and **(E-F)** day 14 treatment. N=5, mean ± SD. p<0.05. One-way ANOVA with Tukey’s multiple comparison test.

Based on the high TRAP activity observed at 7- and 14-days post-surgery, we hypothesized that delivering TBP-NP_IR780_ during this window would result in higher accumulation and longer retention than treatment at D3. We observed highest accumulation of TBP-NP_IR780_ in the D7 treatment group, where TBP-NP_IR780_ accumulated ∼3-fold compared to SCP-NP_IR780_ (TBP-NP: 3.2 x 10^10^; SCP-NP: 8.4 x 10^9^, p < 0.01) and resulted in sustained retention for 14 days after treatment **(Fig. 3C, D)**. However, in the D14 treatment group, we observed a ∼2-fold lower initial accumulation of TBP-NP_IR780_ relative to SCP-NP_IR780_ (TBP-NP: 1.9 x 10^10^; SCP-NP: 8.8 x 10^9^, p < 0.01) which resulted in relatively rapid clearance and poor retention **(Fig. 3E-F)**. Collectively, these data demonstrate that TBP-NPs substantially enhance targeting of systemic treatments to the healing tendon and establish that delivery on D7 post-surgery results in both high-efficiency targeting and sustained retention of TBP-NPs.

### Tendon-targeting of TBP-NPs is dose-dependent

Although we achieved high homing and retention of TBP-NP_IR780_ in the tendon, we also observed some off-target accumulation in the contralateral tendon (**Fig. S5**). Therefore, we sought to investigate dose-dependent effects of TBP-NPs to minimize off-target tissue accumulation. IVIS imaging and analysis of total radiant efficiency at the right hind paw showed that there were no differences in NP accumulation between SCP-NP_IR780_ and TBP-NP_IR780_ when delivered at a dose of 5 mg/kg (**Fig. 4 A-B, Fig. S6**). Furthermore, both 5mg/kg TBP-NP_IR780_ and SCP-NP_IR780_ were cleared within 7 days, compared to the 50 mg/kg group where TBP-NP_IR780_ were retained for 14 days, and SCP-NP_IR780_ were retained for 7 days.

**Figure 4.**
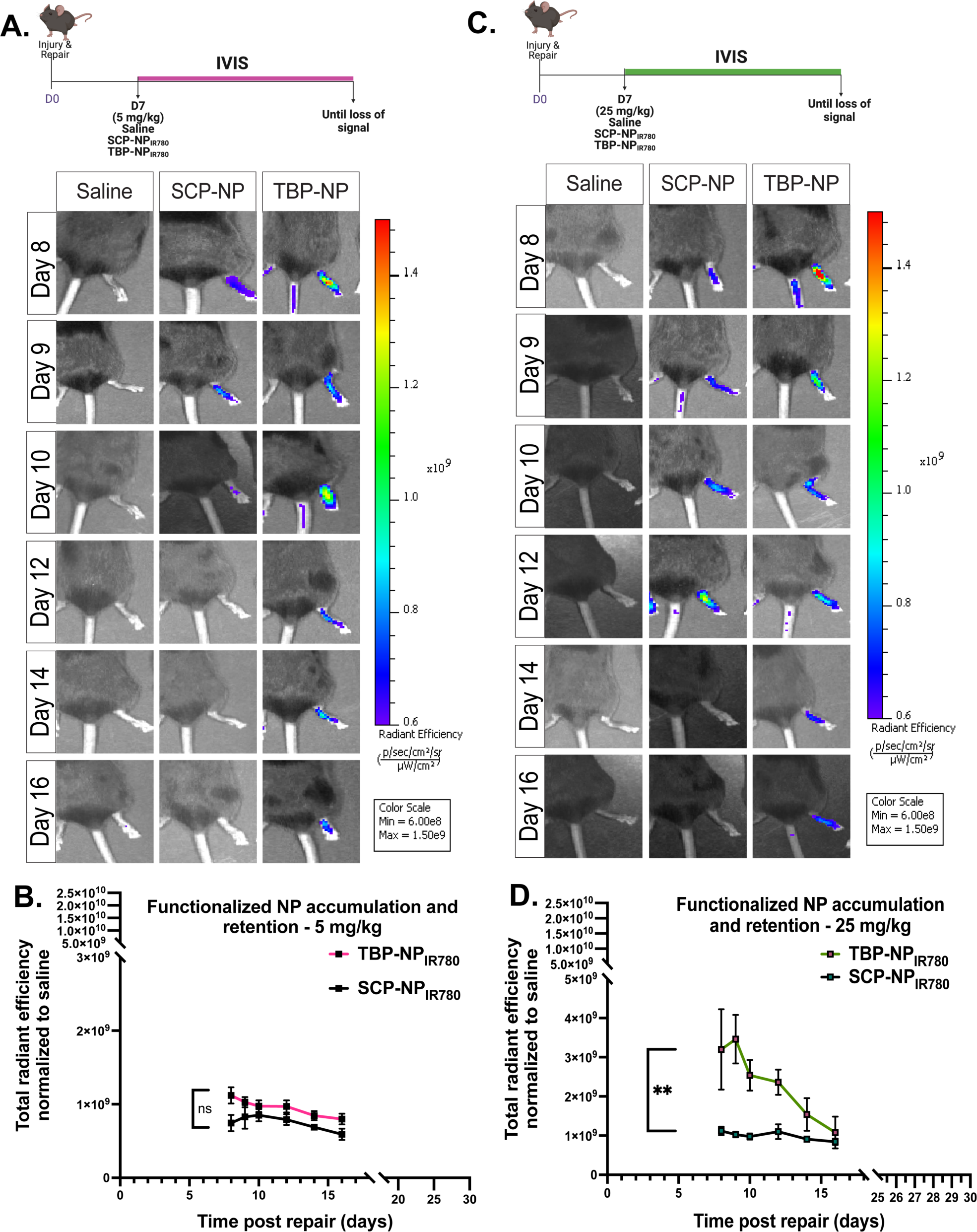
High homing of TBP-NP to the tendon is dose dependent. Schematic showing treatment timeline of TBP-NPs delivered at **(A)** 5mg/kg and Representative IVIS images showing accumulation and retention of 5 mg/kg NPs. **(B)** Graph showing minimal tendon targeting of TBP-NPs versus SCP-NPs and saline control after treatment at 5 mg/kg. **(C)** Schematic showing treatment timeline of TBP-NPs delivered at 25 mg/kg and representative IVIS images showing accumulation and retention of 25 mg/kg NPs. **(D)** Graph showing increased tendon targeting of TBP-NPs versus SCP-NPs and saline control after treatment at 25 mg/kg. N=5, mean ± SD. p<0.05. One-way ANOVA with Tukey’s multiple comparison test.

NPs delivered at a dose of 25 mg/kg resulted in a ∼2-fold higher accumulation of TBP-NP_IR780_ compared to SCP-NP_IR780_. However, the extent of this accumulation (1.9 x 10^10^) was still significantly lower than that observed in the 50 mg/kg treatment group (3.2 x 10^10^, p = 0.01), resulting in faster clearance (10 days vs. 14 days with 50 mg/kg) (**Fig. 4 C-D, Fig. S6).**

### TBP-NPs are biocompatible

Given the promising targeting and retention behavior of the 50 mg/kg TBP-NP_IR780_ administered at D7, this dose and timing regimen was used for all subsequent experiments, and we next assessed systemic biodistribution of the NPs and any potential cytotoxicity. Tissue distribution was analyzed by IVIS 24 h after injection by measuring IR780 signal in organs. While the signal was almost undetectable in several organs (heart, femur, spleen, and CL tendon) in all treatment groups, we observed a considerable IR780 signal in the liver in both TBP-NP and SCP-NP treatment groups (**Fig. 5A-B**). Further quantification of IVIS signal revealed that while maximum radiant efficiency was recorded in lung and repair tendon tissues, lower radiant efficiencies were observed in the liver, spleen, and femur (**Fig. 5C**), suggesting that the high overall accumulation of NPs in the liver was, in part, due to the large liver volume. To further confirm that there was no toxicity after treatment with 50 mg/kg NPs, we investigated the morphology of specific compartments of the liver and kidney. Specifically, we stained liver and kidney sections for hematoxylin and eosin (H&E) to observe structures including the central vein, sinusoids, branch of portal vein, and Kupffer cells in the liver (**Fig. S7A**). Similarly in the kidney, the Bowman’s capsule, Bowman’s space, proximal and distal tubules, capillary lumen, and mesangial cells were evaluated morphologically. There was no difference in the morphology of these structures across all treatment groups, compared to saline, indicating excellent biocompatibility (**Fig. S7A**). Furthermore, we performed an ELISA to investigate ALT and AST enzyme levels. Elevated ALT and AST are associated with liver injury. Compared to saline, we observed no difference in AST (Fig. 5D, Fig. S7B) and ALT (Fig. 5E, Fig. S7B) levels in TBP-NP and SCP-NP groups. Organ-specific spatial distribution of NPs was further characterized via histology and imaging of IR780-labelled NPs. No significant differences in NP area were observed in any tissues other than the healing tendon (**Fig. 5F-G).** More specifically, there was a significant difference in TBP-NP versus SCP-NP uptake in the healing tendon (p < 0.0001) (**Fig. 5G)**, suggesting that targeting of the repair tendon was due to TRAP affinity.

**Figure. 5.**
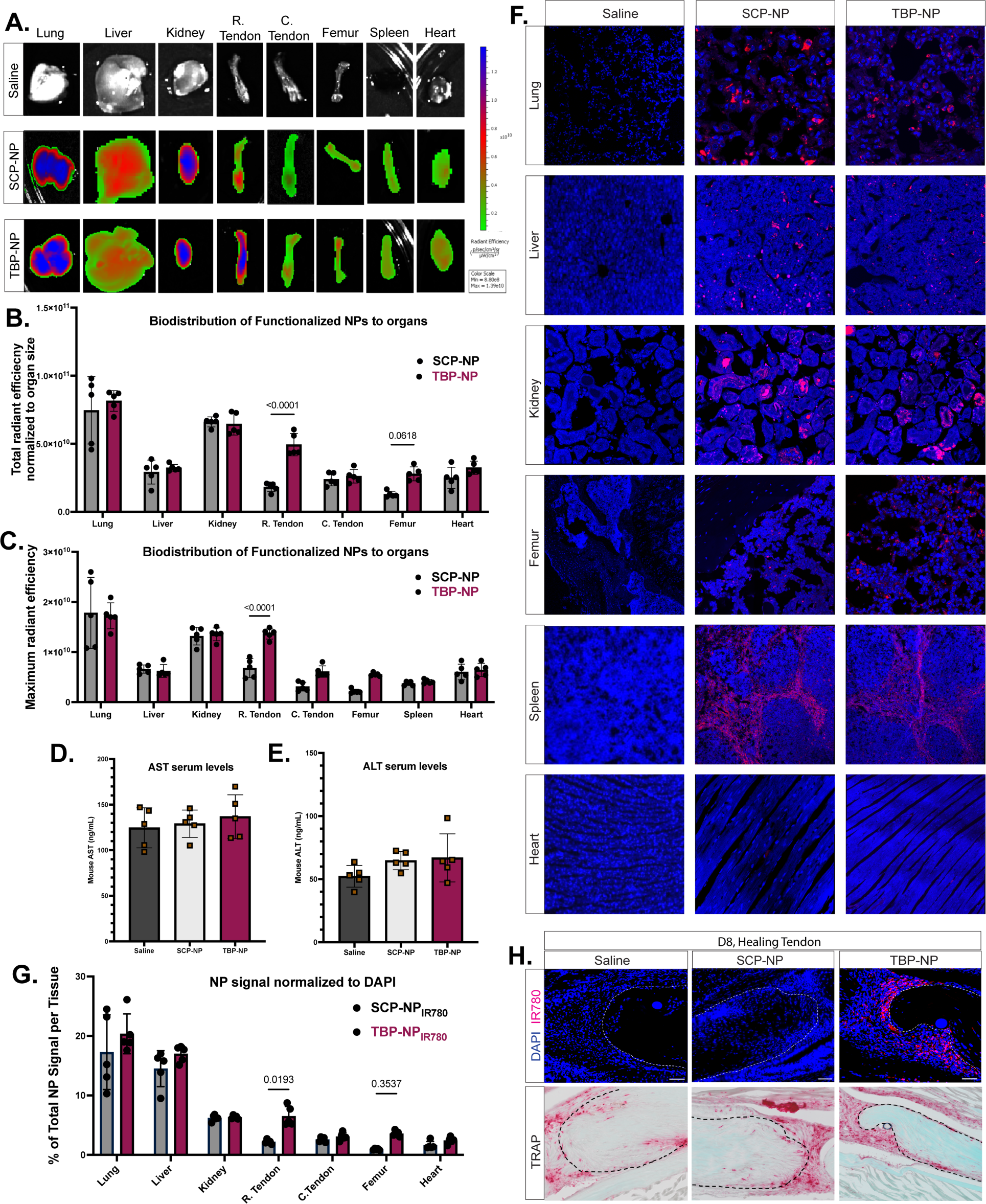
TBP-NP preferentially targets areas of high TRAP activity. **(A)** Representative IVIS images of organs 24 h after treatment with 50 mg/kg nanoparticles on D7. **(B)** Quantification of NP biodistribution in respective organs. **(C)** Quantification of maximum radiant efficiency of NPs in organs after 24 h of treatment. Normal **(D)** AST and **(E)** ALT liver enzyme levels are observed after 24 h of TBP-NP treatment. **(F)** Representative immunofluorescent images showing spatial distribution of NPs within organs after 24 h of treatment (Blue = DAPI, Pink = IR780+ NPs). **(G)** Quantification of NP area within tissues. **(H)** Localization of TBP-NPs within bridging tendon tissue. TBP-NPs are present in areas with high TRAP activity within the tendon. Data was analyzed with 2-way ANOVA with Sidak’s multiple comparison test. N = 5, mean ± SD. P<0.05.

To determine whether areas of high TBP-NP localization were associated with areas of TRAP activity in the healing tendon, tissue sections were stained for TRAP activity following fluorescent imaging. We observed high accumulation of TBP-NPs only within the bridging scar tissue and the periphery of the tendon stubs, which corresponded to areas of high TRAP activity (**Fig. 5H**). However, no SCP-NP accumulation was observed in areas of high TRAP activity (**Fig. 5H**), further supporting the specificity of the TBP approach.

### TBP-NPs within the healing tendon are predominantly internalized by macrophages

Given that many drugs can induce discrete functions in different cell populations, it is critical to understand the cells that take up TBP-NPs. While TRAP activity is typically associated with myeloid lineage cells, including macrophages (**Fig. S8**) and osteoclasts ^29^, some mesenchymal lineage cells can also express TRAP ^30^. Moreover, TBP-NPs increase the homing of NPs to areas of high TRAP activity such that TRAP^-^ cells may also uptake NPs. Therefore, since TRAP is extracellular, we defined the predominant cell population(s) that uptake TBP-NPs using immunohistochemistry and a combination of genetic reporter mouse strains. 24 hours after TBP-NP treatment (**Fig 6A**), approximately 80% of TBP-NPs were taken up by F4/80^+^ macrophages, establishing them as the primary cell population that internalize TBP-NPs (**Fig. 6B-D**).

**Figure 6.**
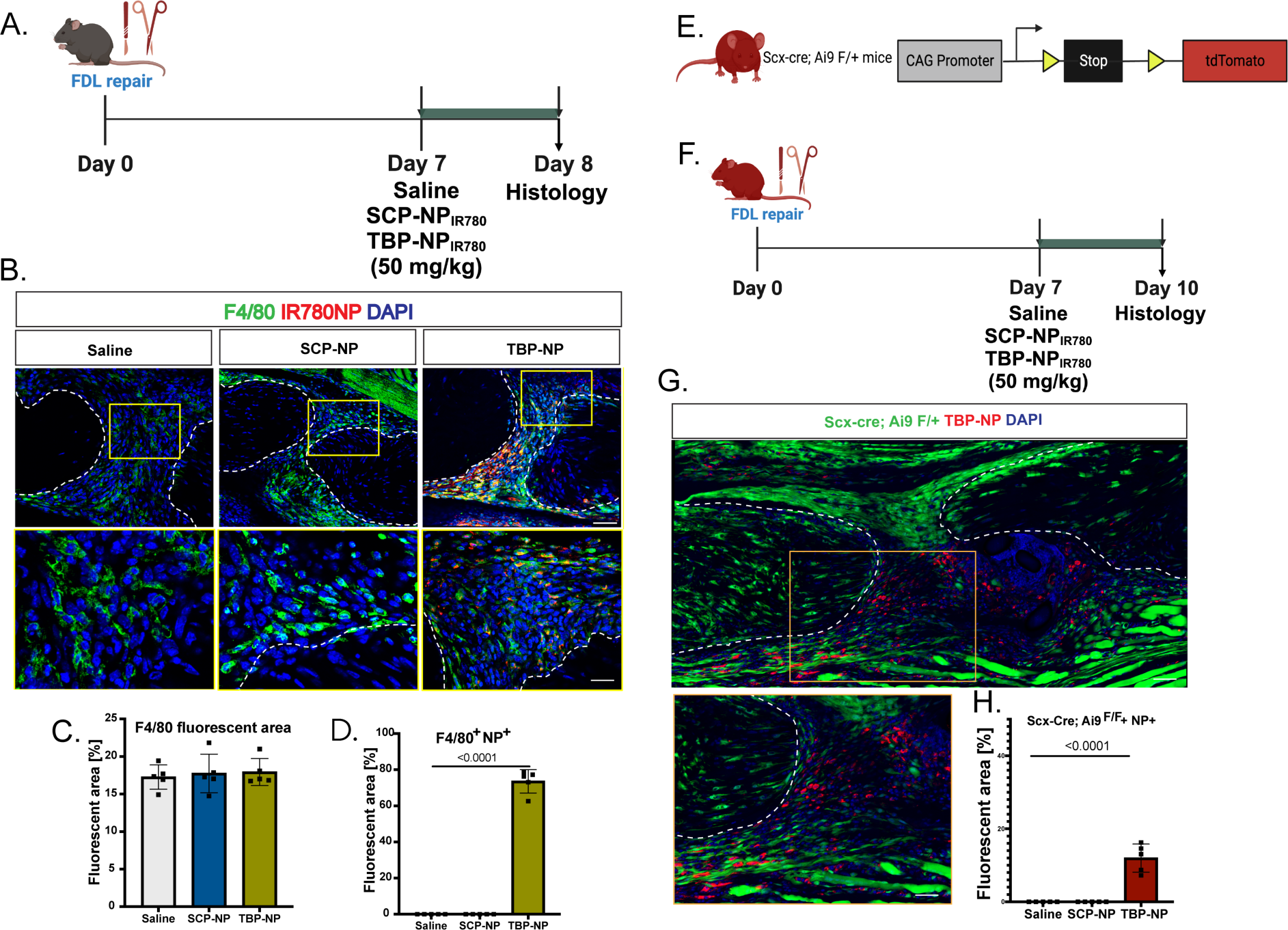
Macrophages are the predominant cell population that internalize TBP-NPs within the tendon. **(A)** Schematic of experimental timeline. **(B)** Immunofluorescent staining and **(C-D)** corresponding quantification showing internalization of TBP-NPs by macrophages within the tendon. **(E-G)** Immunofluorescent staining and corresponding quantification showing expression of S100a4 within the tendon. **(H-J)** Tenocytes, labeled by Scx-Cre: Rosa-Ai9 F/+ expression show limited TBP-NP internalization. **(K)** Quantification of NP+, tenocyte+ cells within the tendon healing environment. Data was analyzed with two-way ANOVA with Tukey’s multiple comparison test. N = 5, mean ± SD.

To determine the extent of tenocyte uptake, Scx-Cre; Rosa-Ai9 mice were used to label tendon fibroblasts (**Fig. 6E-F, Fig. S9**) ^1, 31^. Minimal overlap between TBP-NP_IR780_ and tenocytes (tdTomato) was observed **(Fig. 6 G)** after 72 hours of treatment, with less than 12% of TBP-NPs internalized by tdTomato+ tenocytes (**Fig. 6H**).

### TBP-NPs loaded with Niclosamide inhibit S100a4 expression

The ability to efficiently deliver systemic treatments to the healing tendon represents a major unmet need. While TBP-NPs may be used to deliver a variety of drugs to the healing tendon, we focused on targeted inhibition of S100a4 expression as we have previously shown that genetic knockdown of *S100a4* expression is sufficient to promote enhanced tendon healing^11^. However, this approach is complicated by the limited tissue-specific targeting of systemic treatments ^12^ and poor solubility of drugs, including Niclosamide ^12^. As such, we sought to deliver the S100a4 transcriptional inhibitor, Niclosamide ethanolamine salt (NEN), to the healing tendon via our TBP-NP system. Importantly, the timing of peak TRAP activity precisely overlaps with S100a4 expression in the healing tendon **(Fig. S10)** ^11^ and S100a4 is expressed by both tenocytes and macrophages ^11^.

We achieved efficient loading of NEN into TBP-NPs (75 ± 12 % loading efficiency) as confirmed via HPLC and UV-Vis (**Fig. S11**). No differences in *in vitro* cell viability were observed in the TBP-NP, NEN, and TBP-NP_NEN_ treatment groups compared to saline (**Fig. S12**). To determine whether TBP-NP_NEN_ could inhibit *S100a4* expression in macrophages *in vitro*, BMDMs were treated with either saline, TBP-NP, NEN, or TBP-NP_NEN_ for 48 h. We observed significantly reduced expression of *S100a4* mRNA in TBP-NP_NEN_ and NEN groups, compared to saline (**Fig. S12**). After confirming *in vitro* inhibition of *S100a4* expression, we investigated the ability of NEN and TBP-NP_NEN_ to inhibit *S100a4* expression *in vivo*. While NEN resulted in a significant 44% decrease in *S100a4* expression, relative to saline (p<0.01) (**Fig. 7A**), TBP-NP_NEN_ resulted in a 67% decrease in *S100a4* expression, relative to saline (p < 0.001). Moreover, only TBP-NP_NEN_ treatment resulted in a significant decline in S100a4 protein levels, relative to saline-treated controls (**Fig. 7B, C)**.

**Figure 7.**
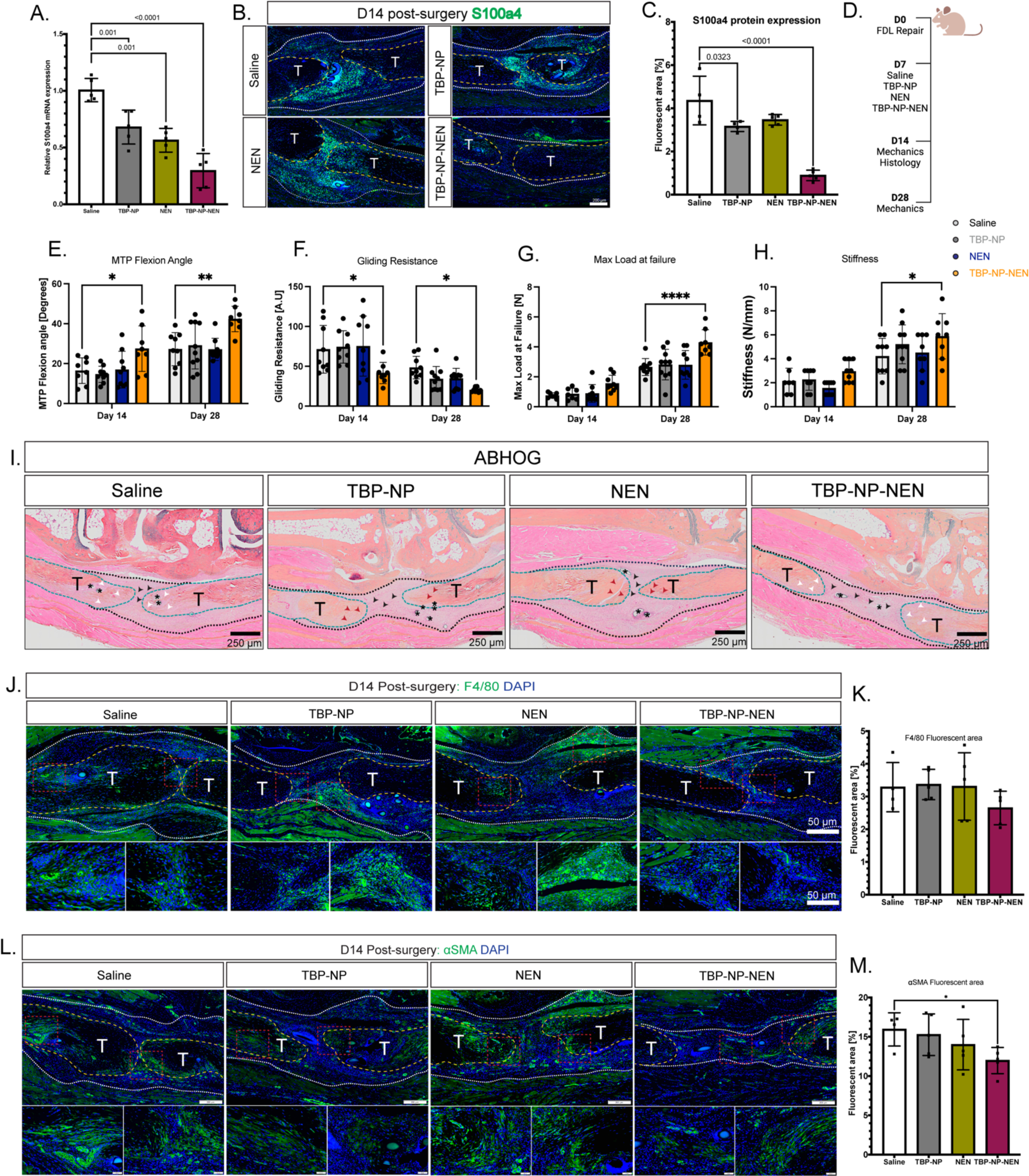
TBP-NP_NEN_ inhibits S100a4 gene expression and promotes regenerative tendon healing. (**A**)TBP-NP_NEN_ inhibits s100a4 gene expression and (**B-C**) Protein expression levels in the tendon. (**D**) Schematic showing experimental timeline. TBP-NP_NEN_ improves mechanical and functional outcomes at D14 and D28 as determined by (**E**) MTP Flexion angle, (**F**) Gliding resistance, (**G**) Max load at failure, and (**H**) Stiffness. TBP-NP_NEN_ treated tendons show improved morphology as shown by (**I**) ABHOG. Tendons demonstrate similar (**J-K**) macrophage content and reduced (**L-M**) myofibroblast content after treatment with TBP-NP_NEN_.

### Tendon-targeted inhibition of S100a4 promotes regenerative tendon healing and improves tendon morphology

Based on our ability to effectively inhibit *S100a4* expression in the healing tendon via the TBP-NP drug delivery system, we assessed the impact of S100a4 pharmacological inhibition on functional and mechanical recovery following injury. Mice were treated with TBP-NP_NEN,_ free NEN, and saline on D7 post-surgery (**Fig 7D**), and tendons were harvested on D14 and D28. No differences in functional measures (MTP flexion angle and gliding resistance) were observed between free NEN, or TBP-NP treated repairs, and saline controls. In contrast, at D14, mice treated with TBP-NP_NEN_ significantly improved gliding function relative to free drug (NEN), TBP-NP, and saline control groups. More specifically, there was a 69% increase in MTP flexion angle (p = 0.03) (**Fig. 7E**), and 42% decrease in gliding resistance (p = 0.001) compared to saline controls. (**Fig. 7F**). These improvements in functional outcomes were maintained at D28, where we observed a 36% increase in MTP flexion angle (p = 0.01) (**Fig. 7E**) and a 58% decrease in gliding resistance (p =0.02) (**Fig. 7F**) relative to saline controls. In terms of mechanical properties, at D14, a 113% increase in max load at failure, and a 37% increase in stiffness was observed in TBP-NP_NEN_ repairs, relative to saline controls, however, these differences were not statistically significant (max load p = 0.08, stiffness: p=0.54). No differences in mechanical properties were observed between saline, TBP-NP, and free NEN at 14 days. (**Fig. 7G-H).** However, at D28, a 62% increase in max load at failure (p < 0.001) (**Fig. 7G**) and 40% increase in stiffness (p = 0.03) (**Fig. 7H**) was observed between TBP-NP_NEN_ and saline controls. These data suggest that TBP-NP_NEN_ treatment improves mechanical and functional outcomes during tendon healing.

### Tendon-targeted S100a4 inhibition modulates macrophage and myofibroblast content

Based on these functional and mechanical changes, we then conducted histological evaluation and profiling of the cell environment during healing in response to TBP-NP_NEN_, relative to free NEN and control groups. Alcian blue Hematoxylin/Orange G (ABH/OG) staining demonstrated deposited collagen fibers bridging the tendon stubs in all treatment groups (**Fig. 7I**, white arrowheads), with a substantial decrease in scar area in the TBP-NP_NEN_ treated repairs, relative to saline, TBP-NP, and NEN controls. Furthermore, the tendon stubs appeared to be actively remodeling in the TBP-NP_NEN_ and saline groups as indicated by the pale orange-pink color observed (**Fig. 7I**, white arrowheads), compared to TBP-NP and free drug controls which displayed the stark orange color of the stubs, characteristic of intact tendon (**Fig. 7I**, red arrowheads). Regarding the cell environment, no changes in the macrophage response (F4/80+ cells) were observed between groups (**Fig. 7J-K**). However, a significant reduction in myofibroblast content was observed in TBP-NP_NEN_ treated repairs, relative to saline controls (28% reduction, p=0.046) (**Fig. 7L-M**).

## Discussion

Developing a tissue-targeting drug delivery system with high specificity, effective drug loading, and robust stability that increases local drug bioavailability is critical for successful translational drug delivery approaches. Poor tissue-specific targeting has limited delivery of promising pharmaco-therapies to the healing tendon. Prior work has demonstrated successful NP-mediated siRNA delivery to enhance tendon healing ^32^. However, local injection was required to ensure tendon localization, and there is some concern that local injections may initiate or re-activate the inflammatory response. Thus, there is a substantial need to facilitate the recruitment of systemic treatments to the tendon repair site. Here, we leveraged a unique spatial transcriptomics dataset to identify a healing niche enriched for *Acp5* expression and TRAP activity. This facilitated the use and refinement of a peptide-functionalized nanoparticle system to enhance tendon-targeted drug delivery.

This nanoparticle drug delivery system is composed of a TRAP binding peptide conjugated to PSMA-b-PS nanoparticles and was previously developed and used by our group in fracture healing applications to expedite bone healing ^19, 33, 34^. In our previous application, carbodiimide chemistry was used as a conjugation method to tether the TBP to the NPs. Although carbodiimide chemistry is a common conjugation method, the maleic anhydride ring opening reaction used in this study provided better control ^22^. The hydrophilic portion of PSMA-*b*-PS contains cyclic anhydrides (MA), which can undergo a nucleophilic addition-elimination reaction when exposed to primary amines. This reaction opens the MA ring and concurrently forms peptide-polymer conjugates, generating carboxylates essential for polymer amphiphilicity ^22^. Remarkably, this process offers a straightforward and robust conjugation technique that obviates the necessity for activators and sidesteps additional purification steps, resulting in the synthesis of TBP-NPs with attractive physicochemical properties and high specificity which facilitated high efficiency tendon targeting after systemic administration.

While recent work has identified NP strategies for tendon-targeting, technical challenges such as the need for local injection or intra-operative delivery limit the translational potential of these approaches. For example, while collagen hybrid peptide-poly(lactic-co-glycolic acid) nanoparticles loaded with Rapamycin (CHP-PLGA-RAPA NPs) ^35^, and mesoporous silica nanoparticles (MSNs)^36^ have been used to target injured tendons, local injection was required. Similarly, nanoparticle/plasmid-laden sutures demonstrate effective targeting of growth factors to the tendon^37^, however, surgical intervention was needed for this delivery approach which may limit the therapeutic treatment window. As such, the development of this TBP-NP drug delivery system that facilitates tendon targeting of systemic treatments addresses an important unmet need.

Given the dynamic nature of the tendon healing environment, characterization of the biodistribution of nanoparticles during different phases of healing is essential to understanding if and how drug payloads will be delivered. That is, treating injured tendons at different time points can induce different homing, retention, bioavailability, and even cytotoxicity, due to natural fluctuations in signaling, regulation, enzymatic activity, and cells within the microenvironment ^2^. Thus, defining an optimal treatment window is critical to successfully developing an innovative and efficacious drug delivery system that improves tendon healing by encouraging a more regenerative rather than fibrotic, healing cascade. In addition, by defining the optimal therapeutic window for this DDS, we can mitigate the unwanted side effects that typically accompany the high doses or multiple doses to achieve concomitantly high drug accumulation in our tissue of interest. Similar to considerations of treatment timing, treatment dose was an important consideration in this study. Surprisingly, the 25 mg/kg TBP-NP dose resulted in relatively poor performance. The biodistribution and pharmacokinetics of NPs in the bloodstream can change when delivered at different doses due to several factors influencing in vivo behavior. One of these factors includes the clearance mechanisms in the body. The liver is the largest organ in the reticuloendothelial system (RES) and hence accumulates a significant portion of untargeted NPs ^38^. While the interaction between Kupffer cell uptake and NP uptake rates at different doses is not well characterized, recent studies have shown that Kupffer cells take up disproportionately larger amounts of NPs when delivered at lower doses because the NP doses fall below a specific threshold ^33, 39, 40^. It is possible that the 25 mg/kg dose used in this study, regardless of binding affinity for TRAP, fell below this “available binding site threshold”, resulting in faster clearance by the RES and lower retention in the tendon. Further studies which fall beyond the scope of this study are required to properly model the threshold uptake kinetics of our NP drug delivery system and confirm this hypothesis.

While prior work, including our own, has pharmacologically targeted both broad processes (e.g., inflammation), or specific signaling pathways that are hypothesized to regulate fibrotic healing, very few studies have achieved a sufficient level of efficacy to move further along the translational pipeline. Therefore, we wanted to determine the ability of this TBP-NP system to effectively achieve local drug delivery in the healing tendon. Importantly, this system has the flexibility to work with many drug classes, however, here we have focused on inhibition of S100a4 based on our prior work demonstrating improved healing in *S100a4*^+/-^ mice^11^. S100a4 is a small calcium-binding protein which has been implicated in fibrosis in many tissues^9^.

S100a4 can regulate several intracellular pathways, though there is some degree of cell- and tissue-level signaling specificity ^9^. In addition, S100a4 can act via cell-non-autonomous mechanisms through secreted S100a4, which can activate pro-inflammatory pathways, recruit immune cells, and trigger the release of matrix remodeling proteins including matrix metalloproteinases (MMPs) ^9, 10^. In addition, S100a4 is expressed by both macrophages and tenocytes in the healing tendon^11^. This, in conjunction with our data demonstrating that macrophages are the primary cells that uptake TBP-NPs, tendon-targeted inhibition of S100a4 was an attractive target for our initial studies. Finally, while NEN effectively inhibits S100a4, NEN is a small, highly hydrophobic drug (MW = 388.2 D), which results in low solubility and rapid clearance, decreasing the likelihood of efficacy in the tendon. Hence, by loading NEN into the hydrophobic compartment of our NPs, we were able to achieve high solubility, outstanding stability, increased circulation half-life, and higher drug bioavailability, leading to successful inhibition of S100a4 at the tendon repair, and enhanced healing.

This study is not without limitations. While we demonstrate effective inhibition of S100a4 using our drug delivery platform, we have not yet tested the effectiveness of loading different therapeutic candidates across other drug classes. In addition, we have focused on S100a4, partly due to the overlap between peak expression of S100a4 during tendon healing, and the window of peak TBP-NP targeting of the tendon. As such, future work will expand to other drug classes, and therapeutic targeting of processes on either side of this therapeutic window. For example, we will need to determine how effectively the TBP-NP system facilitates therapeutic delivery during the late remodeling phase of healing (∼28 days post-surgery in this model). Furthermore, while we have demonstrated and leveraged the robust TRAP activity in the healing tendon, the functional role of TRAP in the tendon is unclear, and represents an important area for future work.

While this work represents a critical step towards a translationally tenable tendon-targeting drug delivery system, there are several important tasks that remain. More specifically, testing of this drug delivery system in larger pre-clinical models will address scalability, which is an important consideration for subsequent clinical safety and ultimately, efficacy trials. Collectively, this study develops and implements a novel tendon-targeting drug delivery system, and demonstrates the efficacy of the TBP-NP system to delivering promising therapeutics to the tendon and enhance healing to an extent that systemic delivery of free drug alone cannot achieve. Moreover, as this drug delivery system is compatible with systemic administration, there is a high degree of flexibility with treatments applied at multiple points during the tendon healing cascade, thus offering a novel and effective treatment strategy for tendon-related pathologies with broad clinical implications. Finally, given that we have identified tissue enriched with macrophages as the primary site of TRAP activity in the healing tendon, this technology may have broad application to other inflammatory pathologies (e.g., tendinopathy) or tissue fibrosis.

## Materials and Methods

### Study Design

The primary goal of this study was to develop a tendon-targeting drug delivery system to promote regenerative tendon healing. Spatial transcriptomics was used to identify a targeting molecule within the tendon. A drug delivery system, involving a peptide conjugated to a nanoparticle was developed and characterized for in vivo tendon targeting. After characterization, a flexor digitorum longus tendon injury was created in C57BL/6J WT mice, and the nanoparticles which were loaded with a near infrared dye were systemically administered once on specific days during the tendon healing cascade. Live animal imaging and confocal microscopy was used to track the nanoparticles to assess biodistribution, spatial localization, and cellular uptake. Histology, and mechanical testing were used to evaluate healing outcomes after delivering a drug with the established tendon-targeting DDS.

### Polymer synthesis and characterization

All studies utilized distilled/deionized water (ddH_2_O) with a resistivity of at least 18 MΩ or greater. Solvents were 98% pure at minimum. For polymerizations, styrene (STY) (99%, ACS grade) underwent distillation for purification, maleic anhydride (MA) was recrystallized from chloroform, and 2,2’-azo-bis(isobutylnitrile) (AIBN) was recrystallized from methanol. The synthesis of polymer diblocks was carried out according to previously described methods ^19, 22^. In brief, diblock copolymers were synthesized with alternating maleic anhydride (MA) and styrene (STY) in the first block (PSMA) and polystyrene (PS) in the second block. An excess of STY was added relative to MA (4:1 [STY]/[MA]) in the presence of the chain transfer agent (CTA) 4-Cyano-4-dodecylsulfanyltrithiocarbonyl sulfanyl pentanoic acid (DCT) (100:1 [MA]/[CTA]).

DCT was synthesized as previously described ^24^. Briefly, n-Dodecylthiol was added dropwise to a sodium hydride suspension in diethyl ether, followed by carbon disulfide and iodine. After purification steps, a dry intermediate was reacted with 4,4’-azobis(4-cyanopentanoic acid) in ethyl acetate under reflux conditions. The final product was obtained through recrystallization using hexanes ^24^. The radical initiator 2,2′-Azo-bis(isobutylnitrile) (AIBN) was used (10:1 [CTA]/[initiator]) in dioxane (57% w/w), and the reaction was purged with nitrogen for 45 minutes before reacting in a 60 °C oil bath for 72 hours.

Subsequently, reactions were exposed to air and dissolved in acetone, followed by precipitation in petroleum ether. The precipitated polymers were then dried under vacuum overnight. For characterization, number average molecular weight (Mn), weight average molecular weight (Mw), and polydispersity (Ð) were determined using gel permeation chromatography (GPC). The mobile phase used for GPC measurements was DMF with 0.05 M LiCl. GPC measurements were performed on a Shimadzu 20A GPC system equipped with a TSKgel SuperHM-N and complementary guard columns from Tosoh Bioscience. The column was operated at 60 °C, with a flow rate of 0.35 ml/min. Detection was achieved using a miniDAWN TREOS light scattering detector from Wyatt Technology and a T-rEX Refractive Index Detector, also from Wyatt Technology. The data obtained were analyzed using Astra 7.3.2 software. Refractive index increments (dn/dc) of 0.142 mL/g was used to characterize both peptide-functionalized and unfunctionalized polymers as determined previously ^24^.

### Peptide synthesis and characterization

Peptides were synthesized following procedures detailed in our previous studies ^19, 22^. Briefly, a Liberty Blue synthesizer (CEM Corp) was utilized for the synthesis of two peptides: tartrate-resistant acid phosphatase (TRAP) binding peptide (TBP-Alloc) with the sequence TPLSYLK^Alloc^GLVTVG, and scrambled binding peptide (SCP-Alloc) with the sequence VPVGTLSYLK^Alloc^LTG. Fluorenylmethyloxycarbonyl chloride (FMOC)-protected amino acids were used in the synthesis.

The coupling of amino acids was achieved using an activator mix containing 0.5 M O-(benzotriazole-1-yl)-N, N, N’, N’-tetramethyluronium hexafluorophosphate (HBTU) in dimethylformamide (DMF) and an activator base mix containing 2 M N, N – Diisopropylethylamine (DIEA) in 1-methyl-2-pyrrolidinone (NMP). TBP and SCP were synthesized with the lysine group protected with an allylic (acid resistant protection) group to ensure that only the terminal amine could participate in the conjugation reaction. Deprotection of individual amino acids during coupling was accomplished using 5% piperazine in DMF. Following synthesis, the peptides were cleaved from Fmoc-Gly-Wang resin (Millipore, MA) using a mixture containing 92.5% trifluoroacetic acid (TFA), 2.5% H_2_O, 2.5% 3,6-dioxa-1,8-octanedithiol (DODT), and 2.5% triisopropylsilane (TIPS) for 2.5 hours, and then precipitated in ice-cold diethyl ether. The products were subsequently freeze-dried for further analysis. The purity of the synthesized peptides was assessed using high-performance liquid chromatography (HPLC) (Shimadzu LC-20AD HPLC system, SPD-20AV UV-Vis detector) HPLC analysis was performed using a Kromasil C18 column (50 mm X 4.6 mm, 5 µm particle size 100 Å pore size) with a variable wavelength UV-vis detector (Shimadzu) using the following parameters: flow rate = 0.5 ml/min, from 95% to 5% A (0.1% TFA in HPLC-grade water) and 5% to 95% B (HPLC-grade acetonitrile) over 21 mins. Molecular weight was verified using matrix-assisted laser desorption ionization time-of-light mass spectrometry (MALDI-TOF) (Brüker Autoflex III). α-cyano-4-hydroxycinnamic acid (CHCA) was used as MALDI matrix and 1:1 acetonitrile:water with 0.1% TFA was used as solvent. Peptide purity was equal to or greater than 95% for the peptides used in this study.

### Polymer-peptide conjugation and nanoparticle self-assembly

The conjugation of polymers and peptides was achieved using the MA ring opening using nucleophilic addition-elimination ^22^. Dry DMF (20% w/v) was used as the solvent to dissolve both PSMA-*b*-PS and peptides. Subsequently, peptides with anhydride-based feeds of 10% were added to the polymer solution and stirred at room temperature overnight. To remove any unreacted peptide and DMF, the resulting peptide-conjugated PSMA-b-PS was precipitated in diethyl ether twice, after which the product was dried under vacuum overnight.

Following peptide conjugation, the resulting conjugates underwent re-analysis to analyze molecular weight and polydispersity. The number of conjugated peptides per polymer was calculated based on the molar mass differences between PSMA-b-PS and the respective conjugate, as detailed in our previous work ^22^. Subsequently, the Alloc protecting group for the primary amine of lysine was deprotected. Briefly, the peptide-polymer was dissolved in DMF (20% w/w) and purged with nitrogen for 20 mins. Tetrakis (Pd(PPh₃)₄) (0.25 N eq to peptide) was dissolved and DMF and DCM at a ratio of 2:1 DMF:DCM and purged with nitrogen for 20 mins. Then, the purged Tetrakis (Pd(PPh₃)₄) was quickly added to the peptide-polymer solution, after which Phenylsilane (25 eq to peptide) was added. After 20 mins, the peptide-polymer solution was precipitated in diethyl ether. This reaction was repeated a second time to ensure the complete removal of the allylic protecting group. Peptide-polymer conjugates underwent dialysis in a 10 MWCO dialysis tube, followed by lyophilization for characterization. To confirm the expected weights and the elimination of allylic compounds, the deprotected peptide-polymer conjugates were characterized using GPC and ^1^H NMR. ^1^H NMR (Bruker 300 MHz) was performed on samples dissolved in DMSO.

To form nanoparticles (NPs), ddH_2_O was gradually introduced via a syringe pump at a rate of 24 μL/min into a solution of polymers dissolved in DMF at a concentration of 6.7 mg/mL. Subsequently, the NP solutions were dialyzed against water for a period of 3 days using a membrane with a MWCO ranging from 6-8 kDa. Following this, the solutions were filtered through 0.2 μm sterile cellulose acetate filters. Dynamic light scattering (DLS) was employed to measure size and surface charge of the NPs at 0.1 mg/mL in water, utilizing a Malvern Zetasizer (Worcestershire, UK). Additionally, transmission electron microscopy (TEM) was utilized, as previously described ^19^ to examine the size and morphology of the NPs. For biodistribution analysis, IR780, a hydrophobic model drug that is fluorescent in the near-infrared region, was loaded into the NPs. Briefly, IR780 was dissolved in acetone and 11 ml of IR780 solution at a concentration of 0.782 mg/ml was added to 175 ml of NPs at a concentration of 0.64 mg/ml. The IR780-loaded nanoparticles were then concentrated to 10 mg/mL and stored in a dark environment at 4 °C.

### Nanoparticle drug loading and characterization

After synthesis and characterization of TBP-NPs and SCP-NPs, Niclosamide ethanolamine salt (NEN) was loaded. SCP-NPs and TBP-NPs were diluted to 1.7 mg/ml. 1.2 mg of NEN was dissolved in 100 µL of DMSO and then diluted in 400 µL of acetone. Ratio of TBP-NPs to NEN was 10:1. The drug solution was then quickly added in a dropwise manner to the stirring NP solution. The reaction was stirred overnight open to atmosphere to allow evaporation of acetone in a chemical fume hood. After loading, the NPs were centrifuged twice at 3000 RPM for 10 mins each, to remove unloaded drug precipitates. Then the drug-loaded NPs were purified using 6 rounds of centrifugal filtration (4000 RPM for 10 mins) to remove remaining organic solvents, free polymer, and unloaded drug. Drug loaded NPs were then diluted in water to a concentration of 40 µM for characterization via HPLC and UV-Vis. The retention time for the NEN was 12 mins with detection at 333 nm. Drug loading efficiency was calculated as loaded drug mass/NP mass x 100%. To further confirm drug loading efficiency, UV–Vis spectra were acquired using a Shimadzu UV-2401PC/2501PC instrument. Before subjecting the drug-loaded NPs to UV analysis, they were first dissolved in water, as described above. The spectra were collected using 1-ml cuvettes possessing a 10-mm path length, over 700 to 190 nm. Absorbance at a wavelength of 333 nm was used, and data was normalized to a standard curve to determine the concentration of NEN and the extent of drug loading efficiency. For in vivo studies, TBP-NPs were injected at a dose of 50 mg/kg, NEN was injected at a dose of 0.75 mg/kg, and TBP-NP_NEN_ was injected at matching doses to TBP-NP or NEN.

### Animal ethics

All studies were carried out in strict accordance with the recommendations in the Guide for the Care and Use of Laboratory Animals of the National Institutes of Health. All animal procedures were approved by the University Committee on Animal Research (UCAR) at the University of Rochester (# 2014-004E).

### Mice

10–12-week-old, female C57BL/6J mice were purchased from Jackson Laboratories (#000664). To trace Scleraxis positive tenocytes, Scx-Cre; Rosa-Ai9^F/+^ mice were used. Scx-Cre mice (a generous gift from Dr. Ronen Schweitzer) were crossed to ROSA-Ai9 mice (#007909, Jackson Laboratories). The ROSA-Ai9 strain expresses the tdTomato fluorescent reporter following Cre-mediated deletion of a loxP-flanked STOP cassette. Following recombination, all Cre-expressing cells, and their progeny permanently express the RFP variant, tdTomato.

### Flexor tendon injury model

Male and female mice, aged 10-12 weeks, were subjected to complete transection and surgical repair of the flexor digitorum longus (FDL) tendon, following a previously described procedure ^1, 11, 21^. Briefly, the mice received a preoperative subcutaneous injection of 15-20 μg of sustained-release buprenorphine to manage pain during the healing process. Anesthesia was induced using Ketamine (100 mg/kg) and Xylazine (10 mg/kg). After removing hair and sterilizing the surgical site, the FDL tendon was cut at the myotendinous junction (MTJ) to transiently minimize strain on the healing tendon. The FDL in the hind paw was exposed through a skin-deep incision on the posterolateral surface, and the FDL tendon was located and completely transected. The tendon was then repaired using 8-0 suture with a modified Kessler pattern, and the skin was closed with 5-0 suture. Post-operatively, the mice were monitored and provided with analgesics as needed for pain management.

### In vivo biodistribution using IVIS

IR780-loaded NPs were retro-orbitally administered on either day 3, or 7, or 14 following FDL injury, using a 30 G needle while mice were anesthetized with Isoflurane (0.7 mg/kg IR780 basis, 50 mg/kg NP basis, 100 µL, 30 G needle). Mice were monitored daily via XENOGEN/IVIS imaging system (PerkinElmer, 780 nm/820 nm for IR780). Semi-quantification of NP biodistribution from IVIS imaging was achieved by measuring the total radiant efficiency of regions of interest and normalizing to saline. Tissue distribution was also analyzed by IVIS 24 hrs after injection by measuring IR780 signal in dissected organs. Specimens were also embedded and cut using a cryostat and imaged at 20X magnification using a Leica confocal Microscopy system. CellProfiler^TM^ software was used to determine the area of cells (DAPI) and NP-loaded IR780.

### Perfusion and blood collection

Anesthetic overdose was induced via intraperitoneal injection of ketamine (200 mg/kg)/xylazine (20 mg/kg). At the time of anesthetic induction, 500 µL of blood was collected from the submandibular vein using a 23-gauge needle. Once a deep anesthetic plane was achieved, the skin and abdominal muscle layers caudal to the xiphoid process were incised with Mayo scissors and the liver was reflected caudally to expose the diaphragm. The diaphragm was removed to expose the thorax and two cuts were made along the right and left of the rib cage. The rib cage was reflected cranially with hemostats. A 1000 mL bag of 0.9% sterile sodium chloride was attached to extension tubing with a 22-gauge needle attached. The needle was inserted into the left atrium and secured with hemostats. The right auricle of the heart was transected with iris scissors and gravity was used to perfuse 0.9% sterile sodium chloride solution into the mouse. Gravity perfusion was performed for 5-8 minutes until all tissues appeared diffusely pale. Blunt dissection was used to remove target tissues for imaging.

For healing studies in awake mice, blood was collected via the submental method. The submental venipuncture method was performed by first restraining the animal by the scruff near the ears so that its head was immobilized and tilted back to expose the submental region, using a grip sufficient to draw back any loose skin or fatty tissue from the access site yet not so tight as to restrict breathing or blood flow to the region. On the midline, a hair whorl was present on the ventral chin. An aim of 1-2 mm to the left or right of this whorl to identify the puncture location was achieved. Then a 4-or 5-mm mouse phlebotomy lancet (or 24-22G needle with 1-2 mm of the needle tip exposed) was used to puncture the skin. The lancet or needle was inserted perpendicular to the skin and withdrawn in a smooth, firm fashion. Blood was allowed to drip freely into the collection vial without manipulating the puncture site. After collection, each mouse was placed in a clean recovery cage and continuously observed until all bleeding ceased. Mice recovered quickly from this procedure and did not require additional observations once bleeding had ceased. No more than 15% of circulating blood volume was removed at a single time point.

### Histology and immunofluorescence

Hind paws from injured C57Bl/6J mice and Scx-Cre; Rosa-Ai9^F/+^ mice were harvested at days 8, 10, and 14 post-injury for both paraffin and frozen sectioning (n = 5-8 per timepoint). For paraffin sectioning, the hind paws were fixed in 10% neutral buffered formalin (NBF) for 72 hours at room temperature, followed by a two-week decalcification process. Subsequently, the paws were processed and embedded in paraffin. Three-micron sagittal sections were cut, deparaffinized, and rehydrated. Antigen retrieval was performed using 10 mM sodium citrate, 0.05% Tween-20, pH 6.0, at 75 °C for 4 hours. The sections were then blocked in blocking buffer (10% normal donkey serum [NDS; #017-000-121, Jackson ImmunoResearch, West Grove, PA, USA] in PBS + 0.1% Tween-20 [PBST]) for 1 hour at room temperature. All secondary antibody incubations were performed for 1 hour at room temperature. Finally, nuclei were counterstained with Hoechst 33342 (NucBlue™ Live ReadyProbes™ Reagent, Thermo Fisher Scientific, Waltham, MA, USA), and the slides were cover-slipped using ProLong™ mounting medium.

For frozen sectioning, hind paws were stored in 10% NBF for 24h at 4 degrees, after which they were decalcified for 72 hr. Following decalcification, tissues were stored in sucrose for 24 h and then embedded and sectioned. All other tissues including heart, liver, spleen, kidney, lungs, and femur were processed in similar fashion. After processing, 10 µm sagittal sections were cut. The sections were then blocked in blocking buffer (10% normal donkey serum [NDS; #017-000-121, Jackson ImmunoResearch, West Grove, PA, USA] in PBS + 0.1% Tween-20 [PBST]) for 1 hour at room temperature. All secondary antibody incubations were performed for 1 hour at room temperature. Finally, nuclei were counterstained with Hoechst 33342 (NucBlue™ Live ReadyProbes™ Reagent, Thermo Fisher Scientific, Waltham, MA, USA), and the slides were cover-slipped using ProLong™ Diamond mounting medium. Imaging was conducted using a VS120 Virtual Slide Microscope from Olympus (Waltham, MA, USA) and a Leica confocal microscope with a far-red scope to allow visualization of IR780+ nanoparticles. The images were subsequently pseudo-colored in OlyVIA (V3.3, Olympus) to facilitate visualization and accessibility, and analyzed in Cellprofiler^TM^ ^41^ as explained in the supplementary section.

### Measurement of gliding function

After euthanization, the hindlimbs were collected at the knee joint. The medial side of the hindlimb was meticulously dissected to release the flexor digitorum longus (FDL) tendon from both the tarsal tunnel and myotendinous junction. The proximal end of the tendon was then secured between two pieces of tape using cyanoacrylate. To evaluate the flexion function of the metatarsophalangeal (MTP) joint, the distal tendon was loaded with weights varying from 0 to 19 g. The MTP flexion angle was recorded for each weight applied. To assess the gliding function, MTP Flexion Angle and Gliding Resistance were computed using established methods ^11^. A lower MTP Flexion Angle and higher Gliding Resistance indicated impaired gliding function, corresponding to increased scar tissue and peritendinous adhesion formation, as previously described ^1^. After conducting the gliding tests, the FDL was freed from the tarsal tunnel. Subsequently, the proximal end of the tendon and the digits were firmly secured within customized grips that opposed each other on an Instron 8841 uniaxial testing system (Instron Corporation, Norwood, MA). The tendon was subjected to a loading process until it reached the point of failure, at a constant rate of 30 mm per minute.

The resulting load-displacement and stress-strain data were analyzed to determine structural properties (stiffness and max load at failure). The stiffness of the tested sample was determined from the slope of the linear region in the load-displacement curve.

### Statistics

Data are expressed as mean ± standard deviation with sample sizes indicated in figure legends. Differences between groups were compared using unpaired Student’s t-tests or one-way or two-way analysis of variance (ANOVA) with Tukey’s or Sidak’s post-hoc testing, as indicated in figure legends. For in vivo studies, 5−10 mice per condition were used. A p-value ≤ 0.05 was used to define statistical significance. Outlier data points were tested for using the ROUT method, and the Q value set at 1%, however no outliers were identified in any quantitative data sets. Statistical analyses were performed in GraphPad Prism 10 Software.

## Acknowledgments

The authors wish to thank Jeff Fox, Dr. Clyde Overby, Dr. Kaye Verda, and Dr. Bradley Nilsson, for their expert technical assistance.

## Funding

National Institutes of Health (NIH/NIAMS)-R21 AR081063 (AEL and DSWB)

National Institutes of Health-The University of Rochester Clinical and Translational Science Institute Trainee Pilot Award UL1TR002001 (EAS).

Schlumberger Faculty for the Future (EAS).

National Institutes of Health S10RR026542-01 (DSWB)

National Science Foundation CBET1450987 (DSWB)

National Science Foundation DMR2103553 (DSWB)

National Institutes of Health (NIH/NIAMS)-R01 AR077527 (AEL)

National Institutes of Health (NIH/NIAMS)-R01 AR073169 (AEL)

## Author contributions

Conceptualization: AEL, DSWB, EAS

Methodology: EAS, IC, BX, YL, JA, CS, AN, KN

Investigation: EAS

Visualization: EAS

Funding acquisition: AEL, DSWB, EAS

Project administration: AEL, DSWB

Supervision: AEL, DSWB

Writing – original draft: EAS

Writing – review & editing: AEL, DSWB, EAS

## Data and materials availability

All data are available in the main text or the supplementary materials

