## Supplementary File for "Development of a Nanoparticle-Based Tendon-Targeting Drug Delivery System to Pharmacologically Modulate Tendon Healing"

### Materials and Methods

*In vitro Binding affinity assay*

The evaluation of TBP-functionalized NP affinity for TRAP was measured using a fluorescamine assay. In brief, TBP-NPs or SCP-NPs (50 µL) were mixed with varying concentrations of TRAP (50 µL) and added to a 96-well black plate (catalog #82050-784, VWR). This mixture was then incubated at 37 °C for 2 hours in 1x DPBS. For each peptide-NP pair, two control groups were included: TRAP only and peptide-conjugated NPs only. After the incubation period, 5 µL of fluorescamine (F9015, Sigma) dissolved in acetone at a concentration of 3.84 mg/mL was introduced to each well. This solution was allowed to react for 30 minutes at room temperature. Subsequently, the fluorescence intensity within the 96-well plate was measured using a BioTek Cytation 5 plate reader, employing fluorescence endpoint settings with excitation at 382/15 nm and emission at 480/15 nm. The collected data were then analyzed in GraphPad Prism using a non-linear fit, one-site, specific binding model [41]. We observed a Kd of -0.221 and 153μM for SCP-NP and TBP-NPs respectively. SCP-NP had a Bmax and $R^2$ of 121.5 and 0.76 respectively, while TBP-NP had a Bmax and $R^2$ or 258700 and 0.97 respectively.

*Nanoparticle dose-response study*

After tendon repair surgery, mice were injected on D7 post-surgery with NP-loaded IR780 retro-orbitally (100 µL, 30 G needle) IR780 at either 5 mg/kg, 25 mg/kg, or 50 mg/kg of NPs. Mice were monitored daily via XENOGEN/IVIS imaging system (PerkinElmer, 780 nm/820 nm for IR780). Quantification of NP biodistribution from IVIS imaging was achieved by measuring the total radiant efficiency of regions of interest and normalizing to saline treated control animals.

*Bone marrow derived macrophage cell isolation and in vitro studies*

Bone marrow derived macrophages (BMDMs) were isolated from 6–8-week-old C57BL/6 mice and cultured in Dulbecco's Modified Eagle Medium (DMEM, Gibco) supplemented with 10% fetal bovine serum (FBS) and 100 units/mL penicillin-streptomycin (Gibco) at 37 degrees Celsius and 5% $CO_2$. Briefly, bones were harvested and transferred into a petri dish with cold PBS, then crushed in DMEM. The media was then filtered through a 40 µm strainer and spun at 350g for 5 mins to pellet cells. Cells were resuspended in 15 mL DMEM with M-CSF (25 ng/mL) and plated. After 24 h, all non-adherent cells were moved to a new flask at 10 million cell density. Media was changed every 72 h until cells reached 70% confluency, at which point they were trypsinized with 0.25% trypsin-EDTA and plated at cell density of 0.15 million cells per well in a 96 well plate for further studies. For cytotoxicity studies, BMDMs were treated with either saline, TBP-NPs (0.22 mg/mL), NEN (10 μM), or TBP-NP-NEN (0.22 mg/mL, 10 μM) 24 h after cell plating. After 48 h, cell viability was determined using Alamar Blue cytotoxicity assay. Briefly, 13.8 μL of AlamarBlue HS reagent was added to wells and incubated at 37 degrees for 3h, after which a cell plate reader was used to read absorbance at 570, 600.

For S100a4 inhibition studies, cells were plated in 12 well plates and treated as mentioned above. After 48 h of treatment, RNA was isolated for qPCR.

*RNA isolation and qPCR*

For in vitro BMDM studies, 500 μL of TRIzol was added to each well of a 12-well plate containing BMDMs, which were previously treated as described. The cells were then scraped into the solution for lysis using a P1000 pipette tip, and the suspension was collected into a 1.5 mL Eppendorf tube. The collected samples were either subjected to immediate RNA isolation or stored at -20 °C. RNA isolation was performed using the Direct-zol RNA Microprep Kit (Zymo Research #R2062) with column purification, including a DNAse step to eliminate DNA contamination. Subsequently, cDNA was synthesized from 200 ng of RNA using an iScript cDNA synthesis kit (BioRad, Hercules CA). Quantitative PCR was conducted with gene-specific primers (Supp Table 1), and the data were normalized to β-actin (Actb) and the expression in WT samples. The relative fold gene expression was determined using the $2^{-\Delta\Delta Ct}$ method. All experiments were conducted with n=3 biological replicates and n=2 technical replicates per sample. For in vivo PCR studies, tendons were harvested from mice at D10 post-surgery by carefully dissecting the skin on the hind paw and transecting the tendon at the heel and bifurcation into the digits. The tendons were immediately flash frozen in liquid nitrogen. The next day, tendons were thawed and immediately transferred into 500 μL of TRIzol reagent, after which they were homogenized. 3-5 tendons were pooled, and cDNA was generated with 500 ng of RNA.

*Quantification of NP presence in tendon*

To quantify NP tissue association, CellProfiler™ Image Analysis Software[42] was used. To determine colocalization of $IR780^+$ NPs, both the pixel-based methods and the object-based methods were used. Briefly, images were loaded into the software, pre-processed using the illumination pipeline, segmented using *three-class Otsu thresholding* to ensure proper spatial mapping of NPs, and then aligned. For pixel-based colocalization measurements, the *measure correlation* feature was used to compare pixel intensities within each channel to determine their correlation, and the correlation coefficient was determined using the Pearson's method. For object-based colocalization measurements, a *parent-child* relationship was set between channels, and the *Relate Objects* and *Classify Objects* pipeline was used. To determine the extent of NP colocalization in respective tissues, the *Measure Image Area Occupied* module was finally used[42].

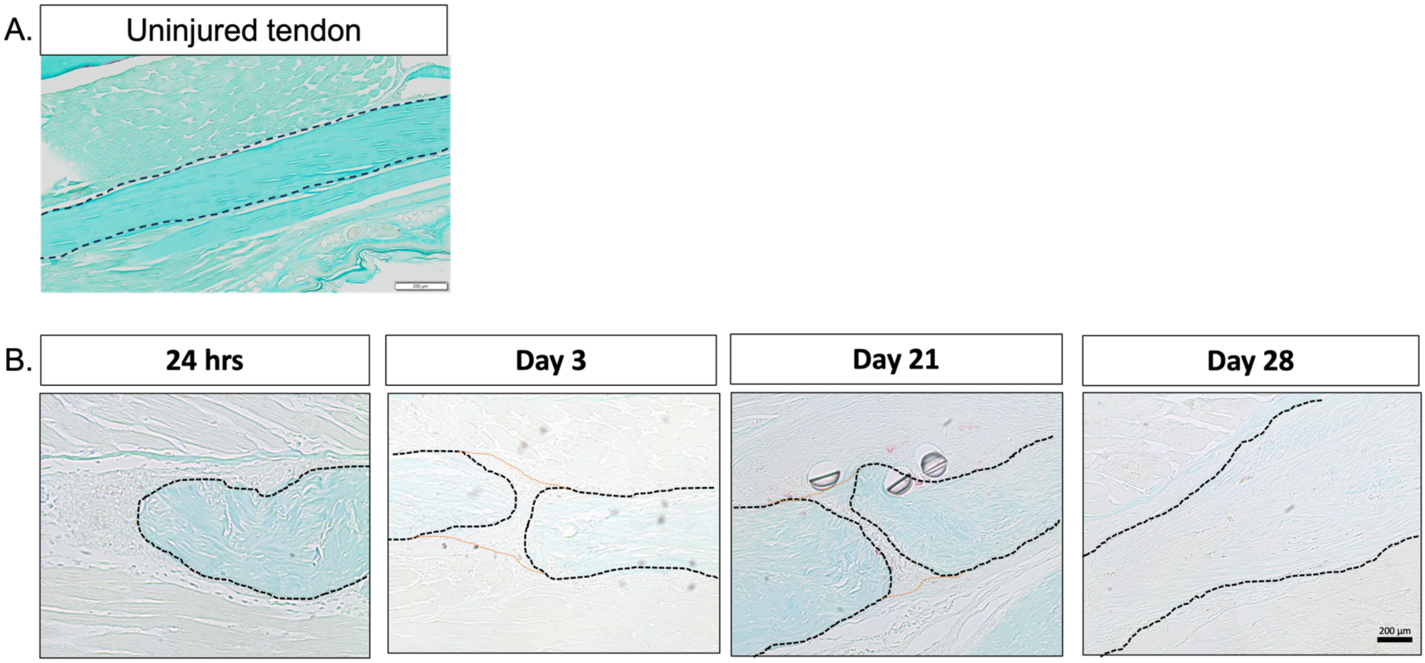


Supplementary Figure 1. TRAP is not present in (A.) uninjured tendons or (B.) within the early inflammatory and remodeling phases of tendon healing.

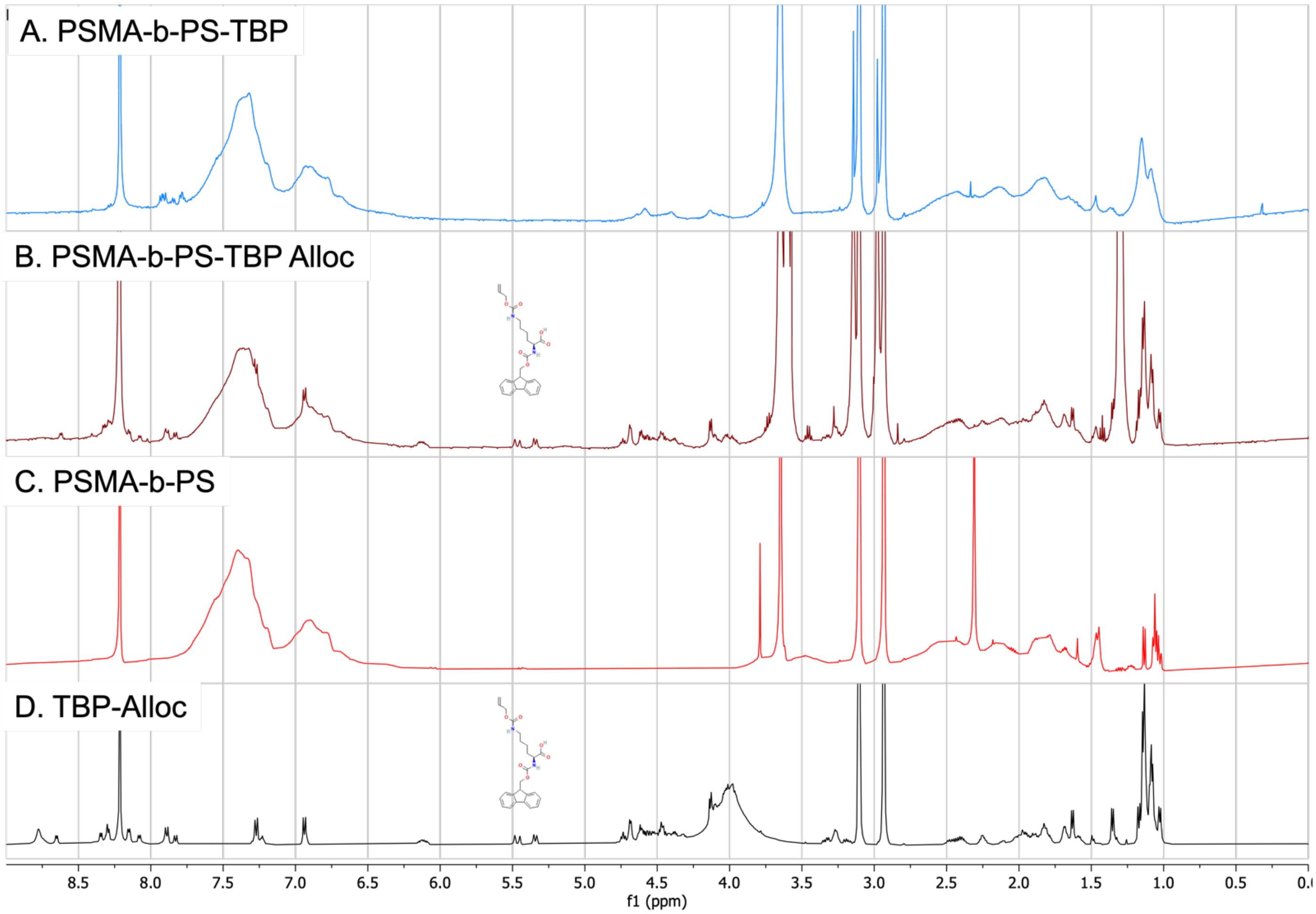


Supplementary Fig 2. NMR confirming absence of allylic peaks. Show structure and peaks.

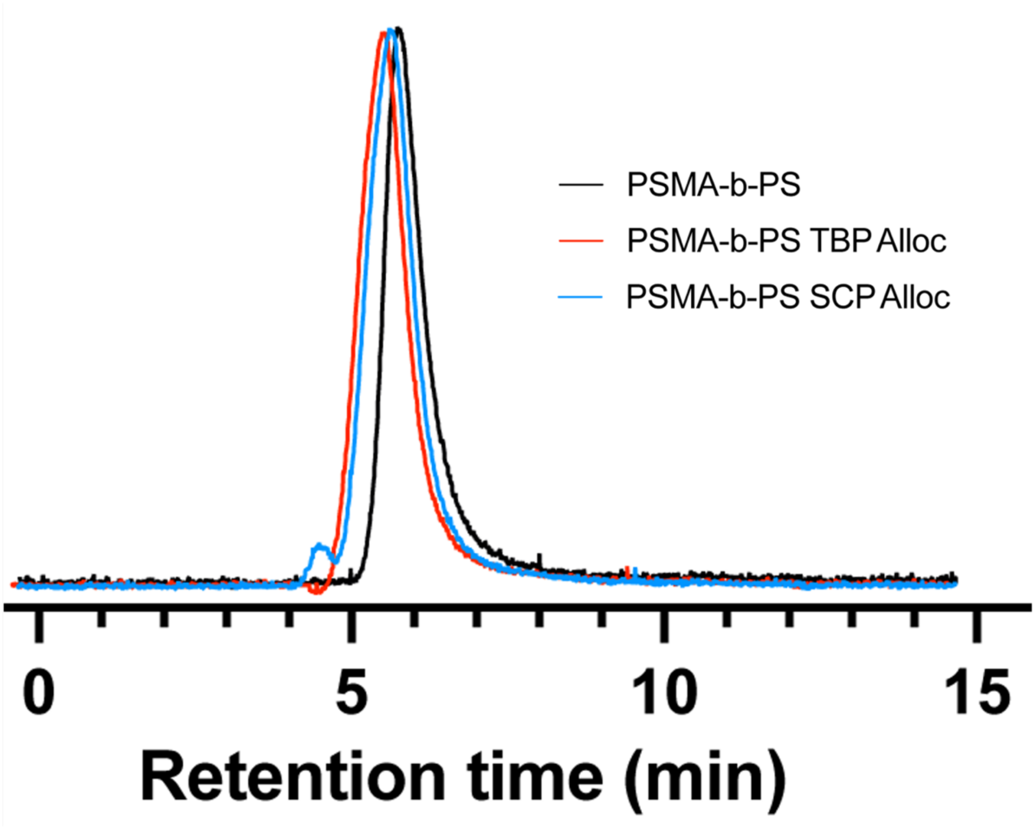


Supplementary Figure 3. GPC chromatograms of unfunctionalized and peptide-functionalized PSMA-b-PS diblock copolymers. Samples were tested at a flow rate of 0.35 ml/min.

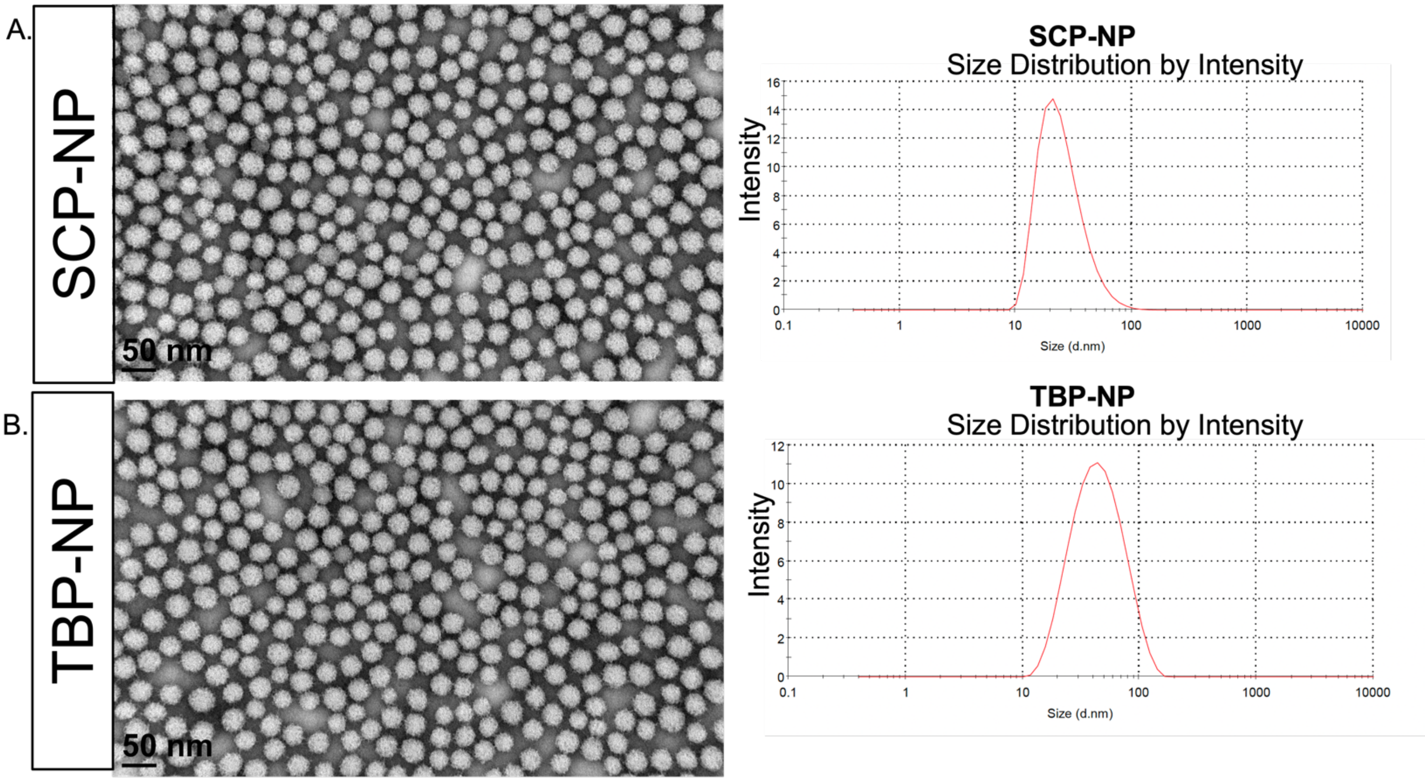


Supplementary Fig 4. TEM and DLS size distribution of NPs. SCP-NP size = ~30nm, and TBP-NP = ~35 nm.

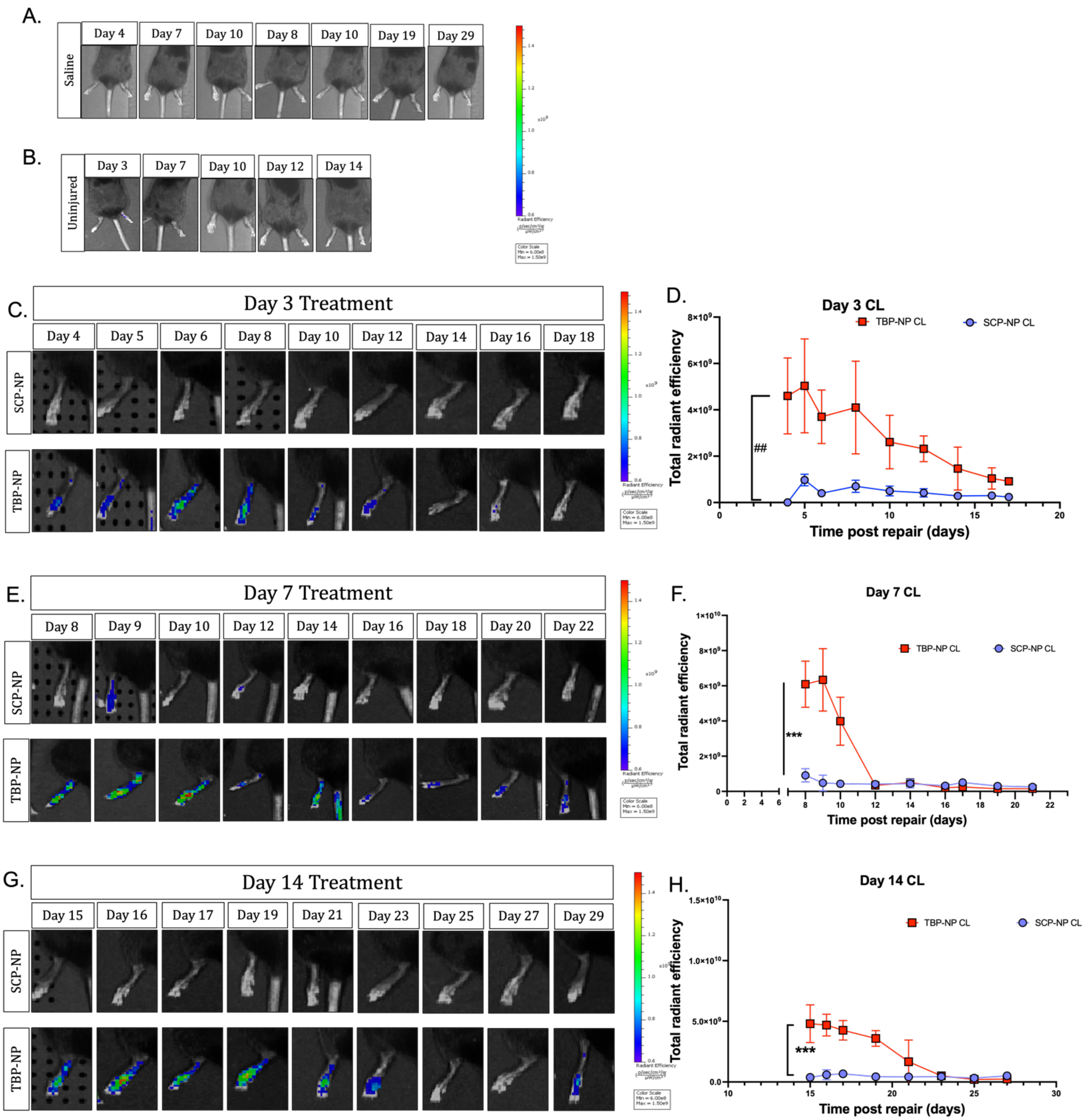


Supplementary Figure 5. Biodistribution of NPs in CL tendon. Minimal off-target effects are observed in contralateral tendons and 50 mg/kg TBP-NP treatment on D7.

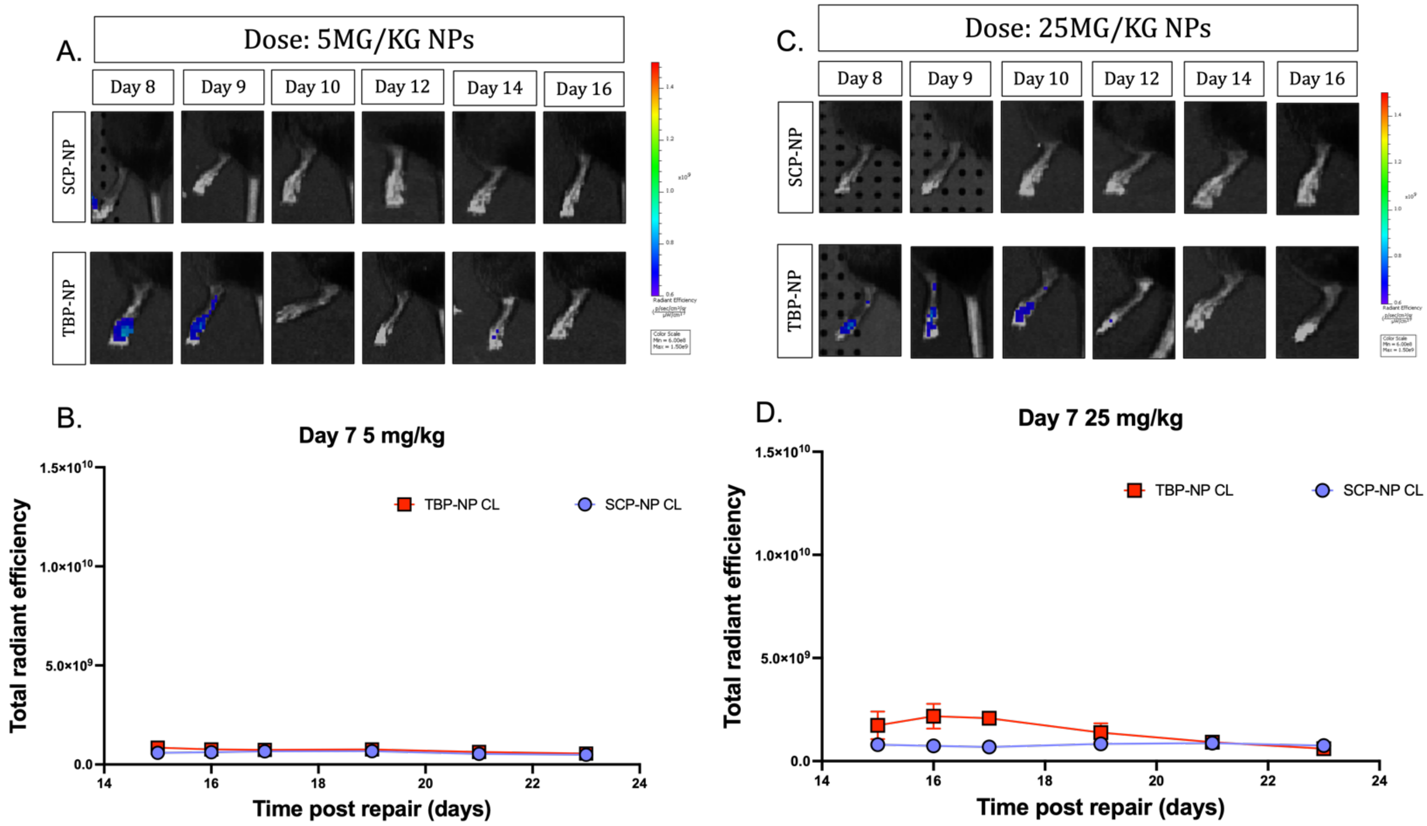


Supplementary Fig 6. Biodistribution of dose-response NPs in CL tendon. No off-target effects are observed in contralateral tendons after 5 mg/kg and 25 mg/kg TBP-NP treatment at D7.

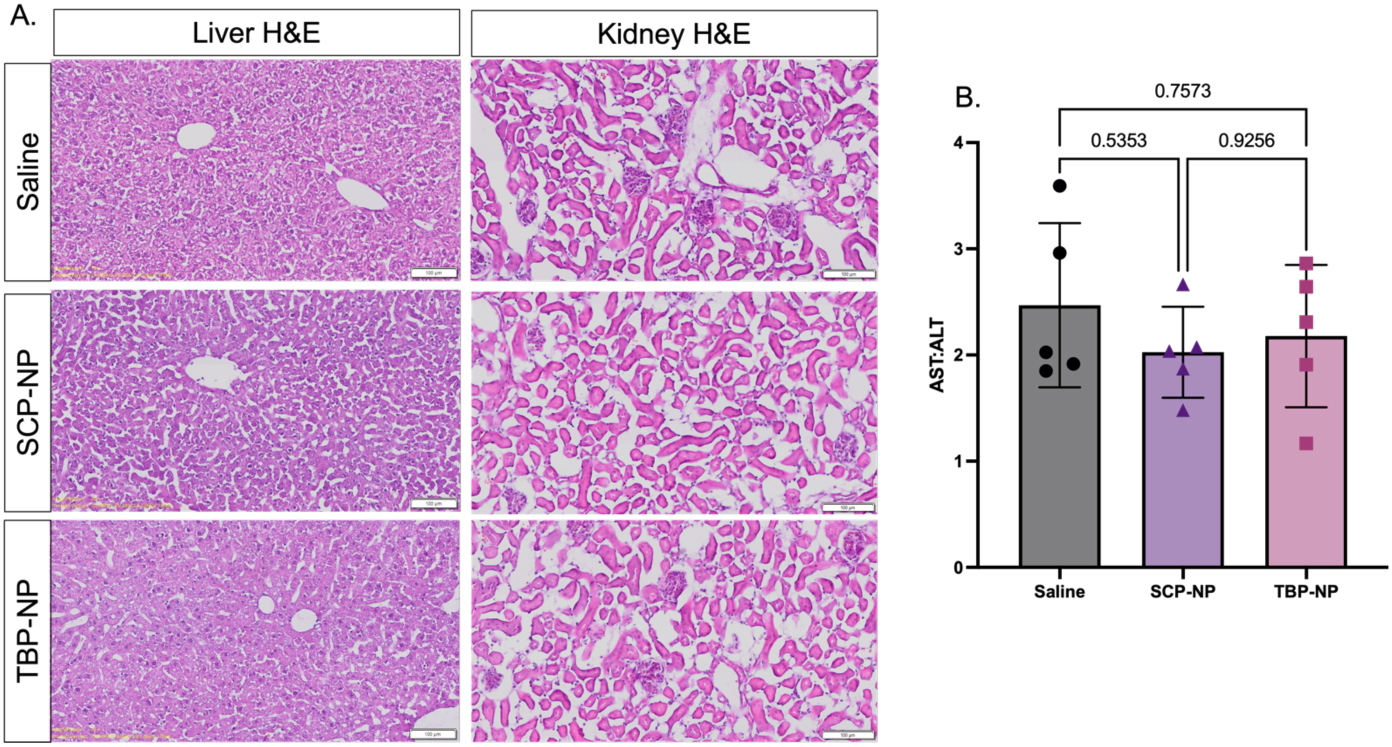


Supplementary Fig 7. Biodistribution of NPs in organs. 50 mg/kg TBP-NPs delivered at D7 post-injury are biocompatible and do not result in liver or kidney injury.

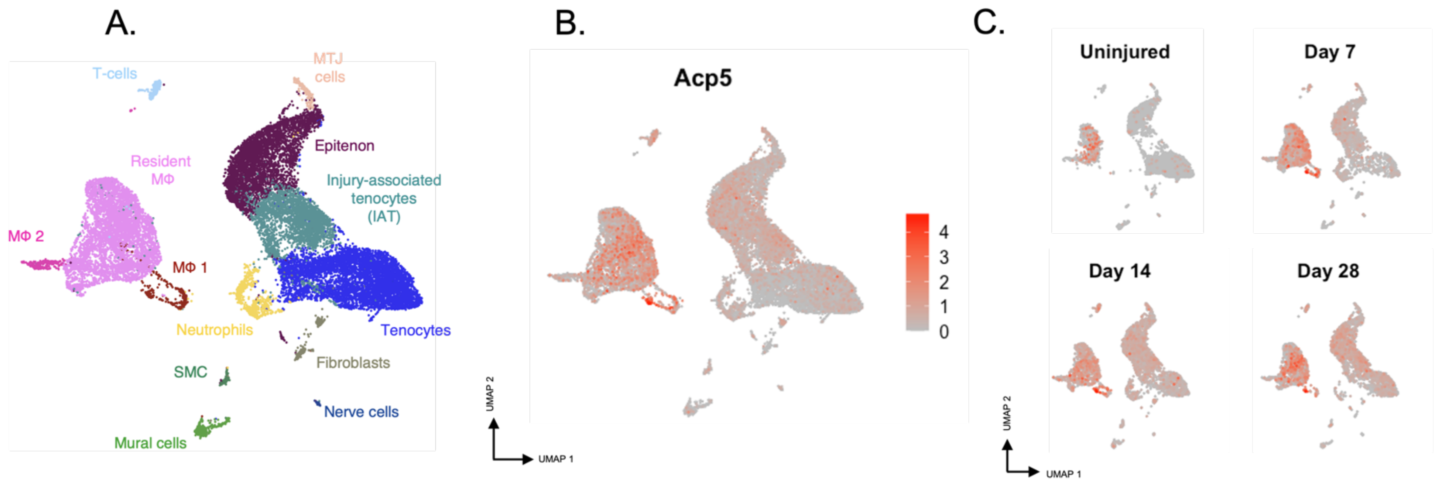


Supplementary Figure 8. scRNA-seq data showing Acp5 expression. Acp5 expression is observed in macrophage clusters.

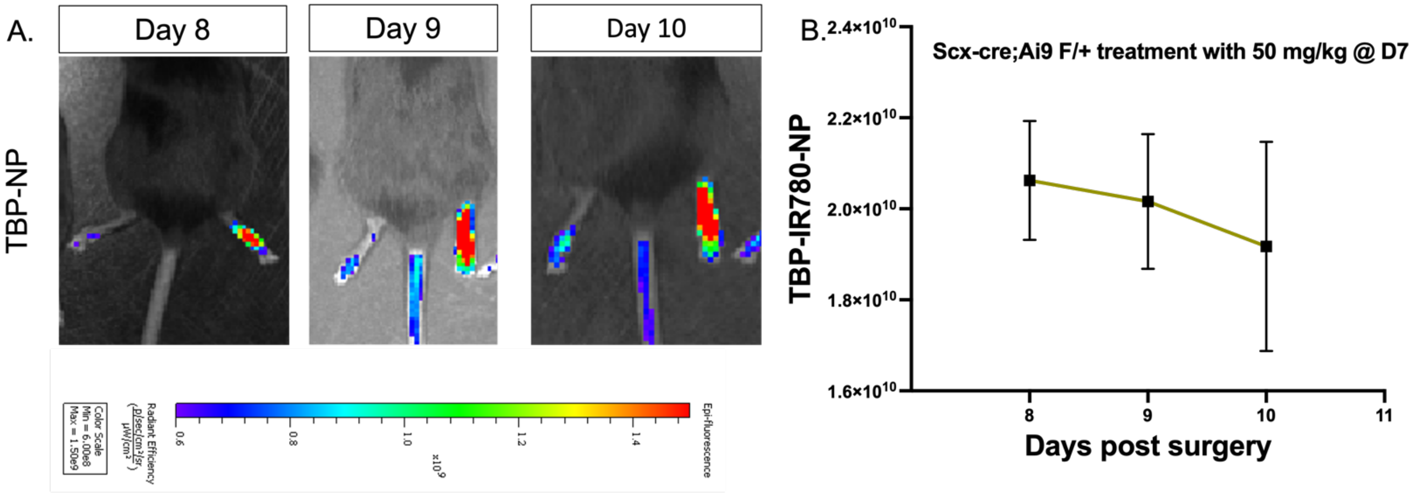


Supplementary Fig 9. Accumulation of TBP-NPs in Scx-cre:Ai9 mice. Similar biodistribution profiles of 50 mg/kg TBP-NPs delivered at D7 post injury are observed in Scx-cre:Ai9 mice, compared to C57B/6J mice.

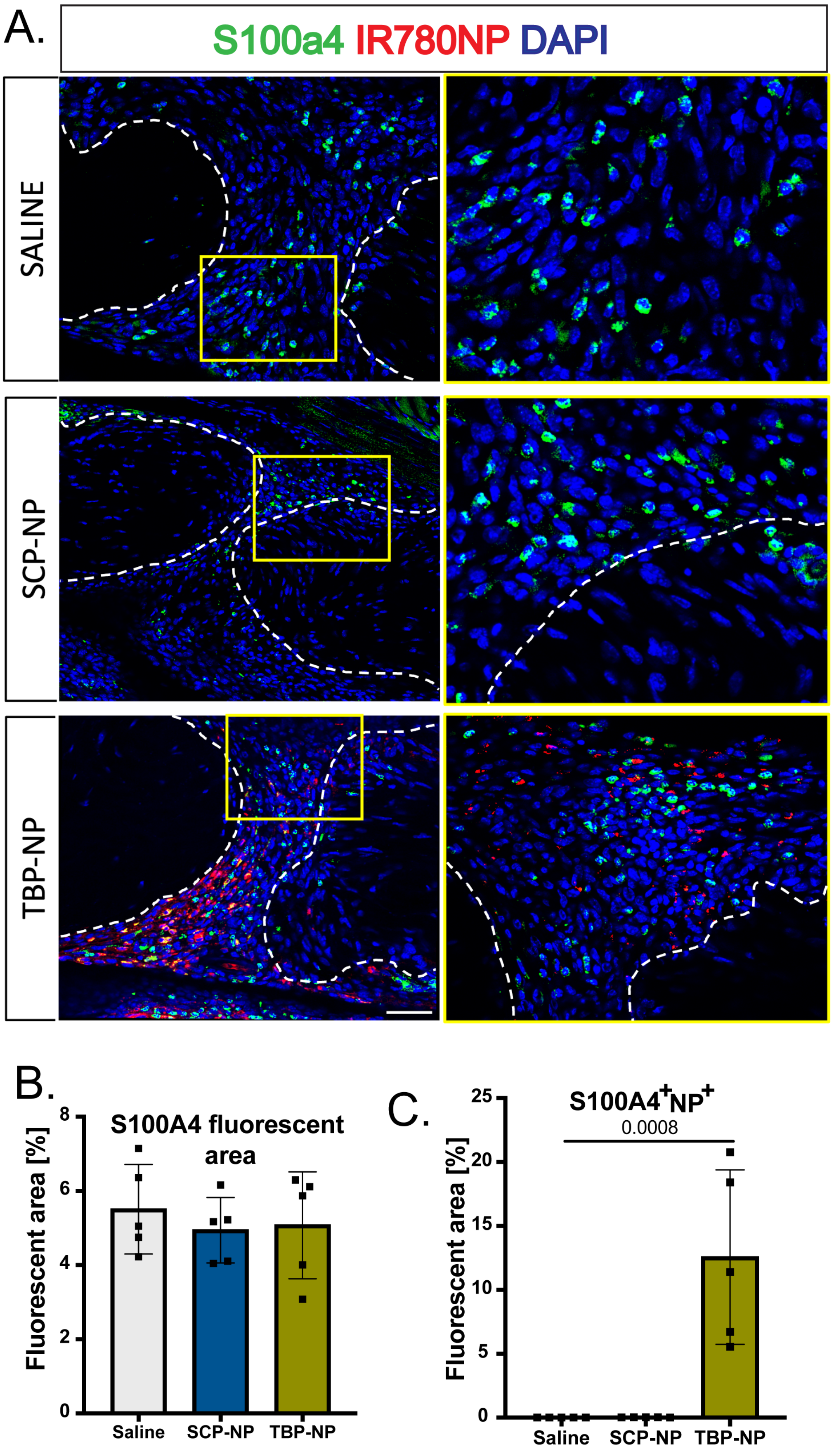
A.
S100a4 IR780NP DAPI
SALINE
SCP-NP
TBP-NP
B.
S100A4 fluorescent area
Fluorescent area [%]
8
6
4
2
0
Saline
SCP-NP
TBP-NP
C.
S100A4+NP+
0.0008
Fluorescent area [%]
25
20
15
10
5
0
Saline
SCP-NP
TBP-NP

Supplementary Figure 10. S100a4 is expressed within the healing tendon during the proliferative phase of healing. TBP-NPs are internalized by a subset of S100a4+ cells.

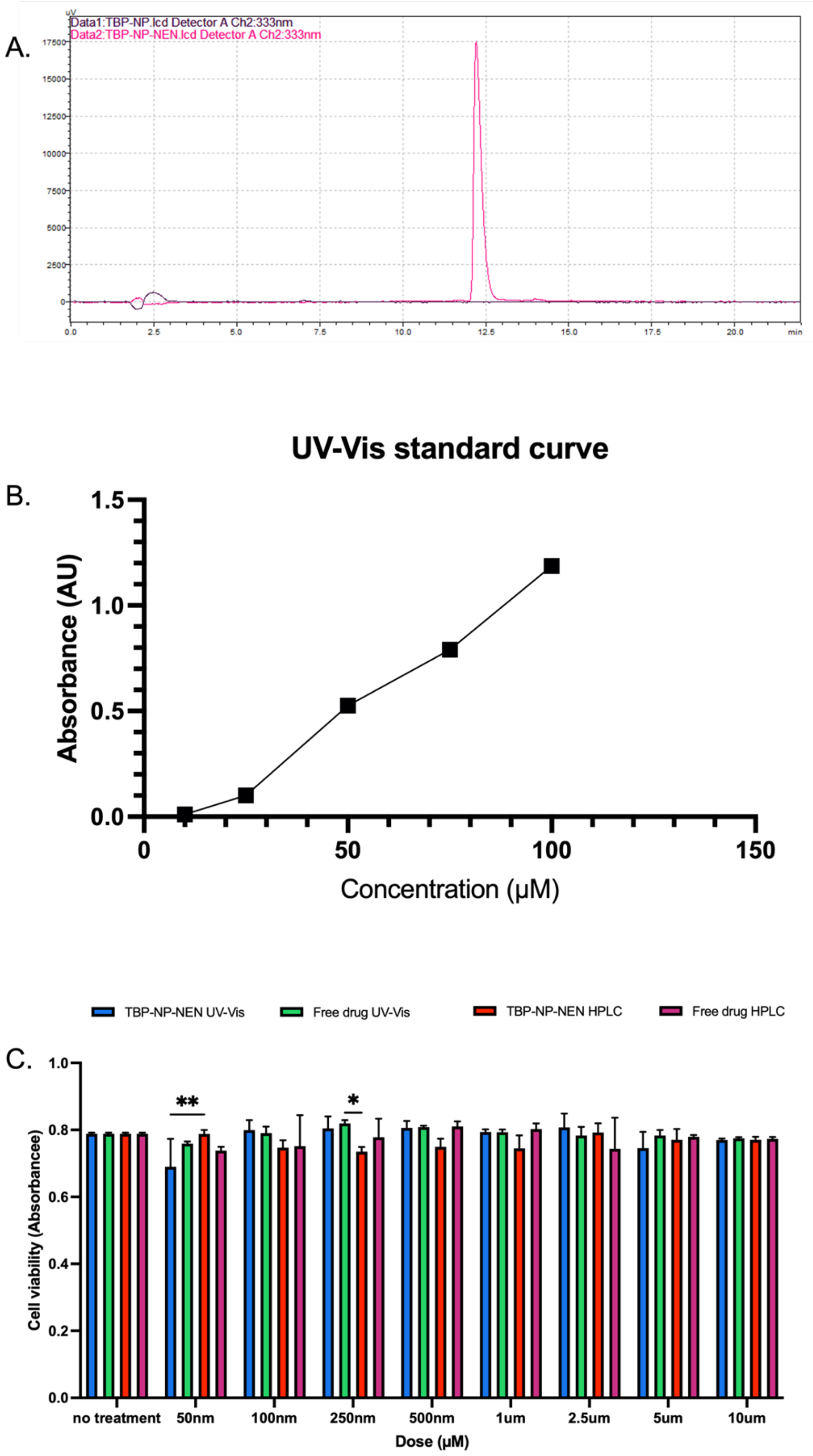

Supplementary Fig 11. Characterization of Niclosamide loading efficiency into TBP-NPs. Both HPLC and UV-Vis were used to characterize Niclosamide loading efficiency, and demonstrate consistent, efficient loading into TBP-NPs. Macrophages treated with TBP-NP$_{NEN}$ have similar viability after LE characterization with both HPLC and UV-Vis. Standard curve wavelength is 333 nm. Wavelength used for HPLC is 333 nm.

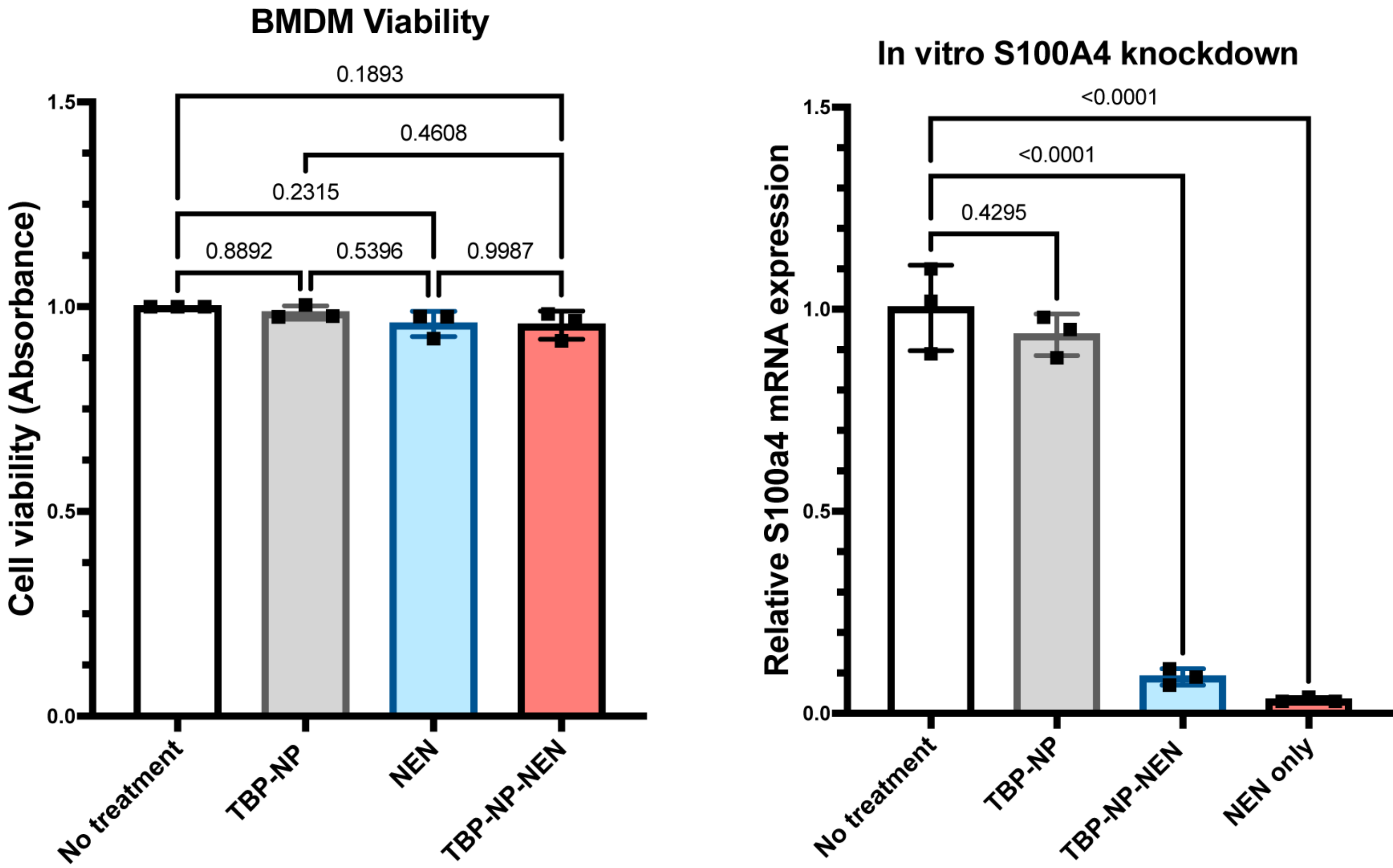


Supplementary Figure 12. TBP-NP$_{NEN}$ is biocompatible. BMDMs treated with TBP-NP$_{NEN}$ maintain excellent viability. TBP-NP$_{NEN}$ efficiently inhibits relative S100a4 mRNA expression in BMDMs after 48h of treatment.
